# Pax3 and Pax7 function in combination with Mitf to generate melanophores and xanthophores in medaka and zebrafish

**DOI:** 10.1101/2023.06.22.546052

**Authors:** Motohiro Miyadai, Hiroyuki Takada, Akiko Shiraishi, Tetsuaki Kimura, Ikuko Watakabe, Hikaru Kobayashi, Yusuke Nagao, Kiyoshi Naruse, Shin-ichi Higashijima, Takashi Shimizu, Robert N. Kelsh, Masahiko Hibi, Hisashi Hashimoto

**Affiliations:** Graduate School of Science, Nagoya University, Nagoya, 464-8602, Japan; Laboratory of Bioresources, National Institute for Basic Biology, Okazaki, 444-8585, Japan; Exploratory Research Center on Life and Living Systems, National Institute for Basic Biology, Okazaki, 444-8787, Japan; Department of Life Sciences, University of Bath, Bath, BA2 7AY, UK

**Author notes:** **Summary statement**: Unique and sequential function of Pax3 and Pax7 allows the diversification of pigment cell types in teleost.

**Keywords:** Paired-type homeobox, Chromatophore, melanocyte, pigmentation, CRISPR/Cas9

## Abstract

Neural crest cells generate numerous derivatives, including pigment cells, and are a model for studying how fate specification from multipotent progenitors is controlled. In mammals, the core gene regulatory network (GRN) for melanocytes (their only pigment cell-type) contains three transcription factors Sox10, Pax3 and Mitf, with the latter considered a master regulator of melanocyte development. In teleosts, which have three to four pigment cell types (melanophores, iridophores and xanthophores, plus leucophores e.g. in medaka), GRNs governing fate specification are poorly understood, although Mitf function is considered conserved. Here, we show that the regulatory relationships between Sox10, Pax3 and Mitf are conserved in zebrafish, but the role for Mitf is more complex than previously emphasised, affecting xanthophore development too. Similarly, medaka Mitf is necessary for melanophore, xanthophore and leucophore formation. Furthermore, expression patterns and mutant phenotypes of *pax3* and *pax7* suggest that Pax3 and Pax7 act sequentially, activating *mitf* expression. Pax7 modulates Mitf function, driving co-expressing cells to differentiate as xanthophores and leucophores rather than melanophores. We propose that pigment cell fate specification is better considered as resulting from the combinatorial activity of Mitf with other transcription factors.

## Introduction

All body pigment cells in vertebrates are considered to derive from neural crest cells (NCCs), a fascinating population of multipotent stem cells (Le Douarin and Kalcheim, 1999; Tang and Bronner, 2020). These cells form an interesting class of NCC derivatives, in part because traditionally all pigment cell-types have been believed to develop from partially-restricted NCC-derived progenitor cells, so called chromatoblasts (Bagnara et al., 1979; Schartl et al., 2016), although we have recently challenged that view, showing that neural crest cells appear to retain much broader multipotency, even in differentiating pigment cells (Kelsh et al., 2021; Subkhankulova et al., 2023). Regardless of their exact nature, these multipotent progenitor cells undergo a process of fate specification to one or more cell-types, before differentiation and eventually commitment to individual specialized cell types. This process is regulated by a complex gene regulatory network (GRN) organised around a core set of transcription factors, which ultimately directs cells to differentiate, stably expressing a transcriptome appropriate to the specific cell-type.

Mammals and birds have only one type of pigment cell, called the melanocyte (Sommer, 2011; Wegner, 2005). The core GRN that controls melanocyte differentiation consists of three transcription factors: a Sry-related HMG-type transcription factor Sox10, a paired-type homeodomain-containing Pax transcription factor Pax3 and a bHLH transcription factor Mitf (Microphthalmia-associated transcription factor). Sox10 and Pax3 cooperatively promote melanocyte development through activating expression of Mitf, which in turn drives melanin synthesizing enzymes, e.g. *Dopachrome tautomerase (Dct)* (Jiao et al., 2004; Lang et al., 2005; Ludwig et al., 2004; Tsukamoto et al., 1992). Mutations in *SOX10, PAX3* or *MITF* similarly cause Waardenburg syndrome in humans (Bondurand et al., 2000; Potterf et al., 2000; Tassabehji et al., 1992), which manifests as patchy pigmentation due to (partial) loss of melanocytes in the skin. Mitf is considered to be a master regulator of melanocytes in mammals as *MITF/Mitf* mutations result in absence of melanocytes, and MITF target genes are numerous and include at least the majority of melanocyte markers (Arnheiter, 2010; Goding and Arnheiter, 2019; Hemesath et al., 1994; Kawakami and Fisher, 2017; Opdecamp et al., 1997; Steingrimsson et al., 1994).

In contrast to mammals and birds, other vertebrates basically have three different types of pigment cell, the melanophore (equivalent to mammalian melanocyte), the iridophore (iridescent cell) and the xanthophore (yellow to red pigment cell). A few species of teleosts, including the Japanese endemic fish medaka, but not zebrafish, have a fourth type, the leucophore (white cell) (Hashimoto, 2021). A more complex GRN must be required to generate this diversity of pigment cell types. Here we address this question of defining the GRN that achieves fate specification and differentiation of the multiple pigment cell in ectothermic vertebrates. We test the hypothesis that the core GRN is conserved in both endothermic and ectothermic vertebrates, while being given added complexity through some additional components specific to ectothermic vertebrates (secondarily lost in the endothermic taxa). Among the three core transcription factors, Sox10 and Mitf have been shown to have highly-conserved roles in melanocyte/melanophore development whereas the function of Pax3 is still poorly understood in other vertebrate species (Greenhill et al., 2011; Hou et al., 2006).

Sox10 plays central roles in all non-ectomesenchymal fates of NCCs, in both fish and mammals, with its loss resulting in absence or severe depletion of all NCC-derived neuronal, glial and pigment cell-types (Britsch et al., 2001; Dutton et al., 2001; Kelsh and Eisen, 2000; Nagao et al., 2018; Southard-Smith et al., 1998). Evidence to date suggests that development of each of these derivatives fails at the earliest stages of specification from the NCCs (Dutton et al., 2001; Elworthy et al., 2003; Elworthy et al., 2005; Greenhill et al., 2011; Kelsh, 2006; Lopes et al., 2008; Petratou et al., 2021; Petratou et al., 2018). Thus, in zebrafish and medaka, loss of Sox10 function results in almost complete lack of melanophores (except for the brain-derived pigmented retinal epithelium), xanthophores and iridophores, while in mice it results in complete lack of melanocytes (Britsch et al., 2001; Southard-Smith et al., 1998).

As in mammals, the zebrafish MITF homologue, Mitfa, plays a crucial role in melanophore development since zebrafish *mitfa^w2/w2^* mutants (also known as *nacre*) lack melanophores throughout life (Lister et al., 1999). A critical role in melanophore fate specification and differentiation is indicated by the extensive absence of melanophore markers in zebrafish *mitfa* mutants and by Mitfa’s activity in directly driving *dct* transcription and melanin pigmentation in the melanophore lineage in zebrafish (Greenhill et al., 2011; Lister et al., 1999) like in mice (Jiao et al., 2004; Ludwig et al., 2004; Tsukamoto et al., 1992). Furthermore, expression of *mitfa* in the NCCs of zebrafish *sox10* mutants is sufficient to rescue full melanophore differentiation (Elworthy et al, 2003), consistent with the master regulator model proposed for mammalian MITF. However, whilst xanthophores are formed in zebrafish *mitfa^w2/w2^* mutants, their development has been noted as abnormal, suggesting that Mitfa may have a partially-redundant role in xanthophore formation (Lister et al., 1999). Nevertheless the exact role of Mitfa in xanthophore development remains unknown.

Pax3 functions as a cooperative partner of Sox10, and thus is also required for melanocyte specification in mammals (Kubic et al., 2008). *Pax3* mutations result in loss of coat color in the *Splotch* mutant mice (Conway et al., 1997; Tremblay et al., 1995), and also in splashed white horses (Hauswirth et al., 2012). In teleosts, however, the role of Pax3 is not clear. An antisense morpholino-mediated knockdown study in zebrafish showed that loss of Pax3 resulted in defective specification of xanthophores but not of melanophores or iridophores (Minchin and Hughes, 2008).

Pax7 is closely related to Pax3 and has been suggested to have similar activity to Pax3. These two transcription factors play important roles in a variety of stem cells, including the NCCs, such as regulating fate decision and differentiation (Buckingham and Relaix, 2015; Nord et al., 2022; Relaix et al., 2004). The similarity is considered to reflect their emergence from a common ancestral gene through duplication (Robson et al., 2006). *Pax7* knockdown by morpholino in the chick indicates that Pax7 is important for induction of the neural crest from the ectoderm, but not specifically required for melanocyte differentiation (Basch et al., 2006). To date, no *PAX7-*associated pigmentation disorders have been reported in humans and mice, although *Pax7*-expressing cells give rise to NCCs (Murdoch et al., 2012). Thus, Pax7 does not appear to play a major role in mammalian nor avian melanocyte development.

Meanwhile, in teleosts, Pax7 seems to be involved in the GRN for pigment cell development. Zebrafish *pax7a; pax7b* double homozygous mutants exhibit complete absence of xanthophores but not of melanophores nor iridophores (Nord et al., 2016), suggesting an essential and specific role of Pax7 in xanthophore formation. However, in medaka, where functional Pax7b has been evolutionarily lost, *pax7a* homozygous mutants exhibit complete absence of xanthophores and leucophores. As a consequence, xanthophores and leucophores have been proposed to share a bipotent progenitor, whose specification requires Pax7a function in medaka (Kimura et al., 2014; Nagao et al., 2014; Nagao et al., 2018). Although that conclusion may need revision in the context of our recent zebrafish observations (Kelsh et al., 2021; Subkhankulova et al., 2023), it remains clear that Pax7 function is necessary for both xanthophore and leucophore development.

Here we used a comparative genetic approach to assess the roles for these key pigment cell transcription factors. We addressed the following key questions: 1) Is Pax3 one of the core transcription factors in the melanophore GRN in medaka and zebrafish; 2) Do Pax3 and Pax7 control specification of other pigment cell types; and 3) Does Mitfa have a role in development of other pigment cell-types in medaka? Our results suggest that Sox10, Pax3 and Mitf have a conserved role as components of the core GRN driving melanophore fate, but that they also work alongside Pax7, to promote xanthophore and leucophore fates, in teleosts.

## Results

### Two paralogous *pax3* genes, *pax3a* and *pax3b*, are identified in medaka and zebrafish genomes

Vertebrate *pax3* and *pax7* are descendants of a common ancestral gene (named *pax37* in ascidians), resulting from the second-round whole genome duplication in vertebrate evolution (Kusakabe and Kuratani, 2007; Wada et al., 1996). In teleosts, two paralogues for each of *pax3* and *pax7* were generated as a result of another round of whole genome duplication (Amores et al., 1998; Postlethwait et al., 2004). Medaka and zebrafish have *pax3a*, *pax3b*, *pax7a* and *pax7b* in their genomes, although medaka *pax7b* lacks three exons (exons 6-8) and appears to be a non-functional pseudogene (See Suppl. Fig. 1).

### *pax3* is expressed in neural crest cells prior to *pax7*

We observed expression of only *pax3b* and *pax7a* mRNAs in medaka NCCs. In situ analyses showed that *pax3b*-expressing cells were observed in dorsal parts of the midbrain, hindbrain and spinal cord, dorsal somites and premigratory NCCs at early somite stages (Fig. 1A-A’’). In contrast, *pax7a* expression was detected in both premigratory and laterally migrating NCCs (Fig. 1 B-B’’). At mid-somite stages, *pax3b*-expressing NCCs were seen in the posterior premigratory region but not in the lateral migratory region, while *pax7a*-expressing NCCs were observed in both premigratory and migrating regions (Fig. 1 C, C’, D, D’, D”). By late somite stages, *pax3b* expression was no longer detectable in the NCCs, but *pax7a* expression was still detected in migratory NCCs (Fig. 1 E, E’) and maintained until almost the hatching stage (Watakabe et al., 2018). These results indicate that expression of *pax3b* in medaka is transient, restricted to premigratory NCCs, and precedes that of *pax7a*, which is maintained into migrating NCCs. These expression patterns are similar to those previously reported for zebrafish *pax3* (*pax3a* and *pax3b*) and *pax7* (*pax7a* and *pax7b*) (Minchin and Hughes, 2008).

**Fig. 1.**
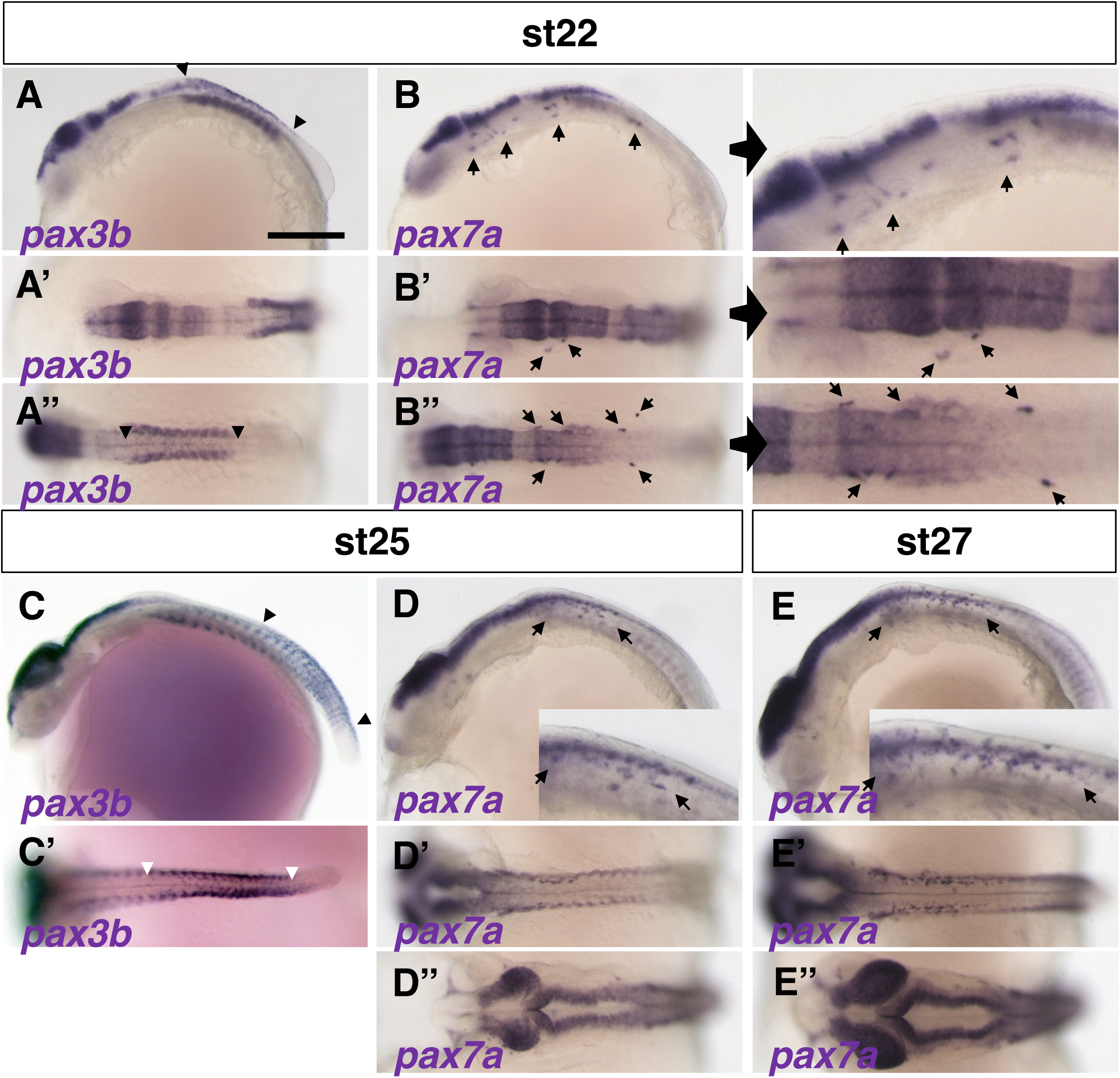
Expression patterns of *pax3* and *pax7* in medaka. (A, C) *pax3b*. (B, D, E) *pax7a*. Expression of *pax3b* mRNA is detected in NCCs at stage 22 (9 somite embryo) (A, A’, A’’, between arrowheads) while *pax7a* expression is observed in migrating NCCs anteroventral to the *pax3b*-expressing premigratory NCCs (B, B’, B’’ and their close-up views on the right panels, *pax7a*-expressing cells are indicated by arrows). The *pax3b* expression has shifted posteriorly at stage 25 (18-19 somite embryo) (C, C’, between arrowheads) and become undetectable by stage 27 (24 somite embryo) (not shown). *pax7a* mRNA is continuously observed in migrating NCCs from stage 25 to stage 27 (indicated by arrows. See also Suppl. Fig. 2). *pax3b* and *pax7a* mRNAs are expressed in similar regions of neural tissue and somites (A-E). The expression of *pax3b* is observed in several anterior somites at stage 22 (A), shifts to posterior somites at stage 25 (C. See also Suppl. Fig. 2), and appears to precede that of *pax7a* in somites (B, D). The time windows of *pax3b* and *pax7a* expression overlap during stages 22-25. (A-E) Lateral views. (A’-E’, A’’, B’’, D’’, E’’) Dorsal views. Scale bars = 250 µm

The above results were confirmed by observing fluorescently-labelled *pax3b*– and *pax7a*– expressing cells in a newly established stable transgenic medaka Tg*(pax3b-hs:GFP)* and previously reported Tg*(pax7a-hs:GFP)* lines (Suppl. Fig. 2 A) (Watakabe et al., 2018). The *pax7a*-GFP remained detectable in NCCs until a late somite stage, much longer than the *pax3b*-GFP (Suppl. Fig. 2 B-H). We also tested whether the expression of *pax3b* and *pax7a* overlapped, by using Tg(*pax3b*-*hs*:*GFP*) crossed with previously reported Tg*BAC*(*pax7a*:*DsRed*) (Nagao et al., 2018). A portion of the cells positive for *pax3b*-GFP, but not all, were also positive for *pax7a-*DsRed (Suppl. Fig. 2 I-K), indicating that these cells express both Pax3b and Pax7a. There were *pax3b*-GFP positive but *pax7a*-DsRed negative cells, suggesting that *pax3b*-expressing cells give rise to other pigment cell population than *pax7a*-expressing xanthophore/leucophore progenitors.

### Loss of *pax3* function results in delayed formation of xanthophores in zebrafish and medaka, and also of leucophores in medaka

To investigate the role of *pax3* and *pax7*, we generated mutations in *pax3a* and *pax3b* in medaka and *pax3a*, *pax3b*, *pax7a* and *pax7b* in zebrafish using TALEN or CRISPR/Cas9 (Supp. Figs. 1, 3). In addition, we used medaka *leucophore free-2 (lf-2)* as a *pax7a* mutant, which was previously reported as a loss-of-function mutation (Suppl. Fig. 1) (Kimura et al., 2014).

We examined pigment cell phenotypes of the mutants. The medaka *pax3b* homozygous mutants showed a delay in the formation of xanthophores and leucophores, which was detected as a reduction in the number of these cell types in hatchlings (Fig. 2 A-D, G, H, Suppl. Fig. 4). This phenotype was much milder than that of the *pax7a* mutant, in which these two cell types are completely lost (Fig. 2 E, F, Suppl. Fig. 4) (Kimura et al., 2014). Medaka *pax3b* mutant fish became apparently normal in pigmentation as they grew, with xanthophores and leucophores restored (Suppl. Fig. 4).

**Fig. 2.**
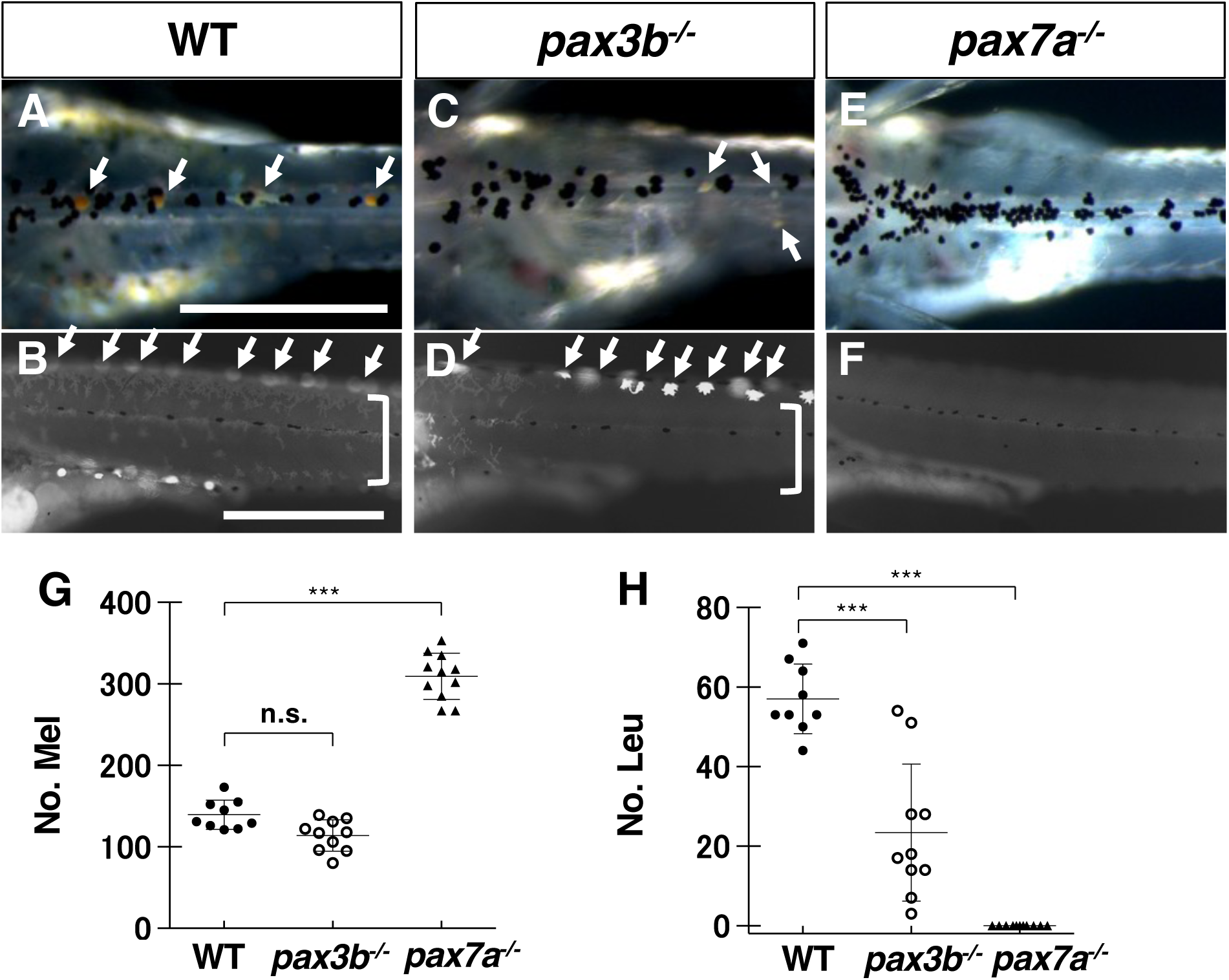
Phenotypes of medaka *pax3* and *pax7* mutants. (A-F) Medaka 9 dpf hatchlings. Pigment cell phenotypes are compared between WT (A, B), *pax3b*^-/-^ (C, D) and *pax7a^-/-^* (*lf-2*) (E, F). (A, C, E) Dorsal views of the trunk in dark field. (B, D, F) Lateral views of the trunk under UV light. (G, H) Cell counts. Melanophores (G) and leucophores (H) on the dorsal surface of the trunk are quantified. Xanthophores and leucophores are severely reduced in number in medaka *pax3b*^-/-^ mutant hatchlings (leucophores indicated by arrows in A, B, C, and D; xanthophores, which are autofluorescent under UV light, indicated by square brackets in B and D), and are completely absent in medaka *pax7a^-/-^* mutants (E, F, H). The number of melanophores is unaltered in *pax3b*^-/-^ mutants (A, C, G), whereas significantly increased in *pax7a^-/-^* mutants (E, G). Significant difference was determined by Kruscal-Wallis test. \*\*\**p*<0.05. n. s. = not significant.

Similarly, zebrafish *pax3a* homozygous mutant hatchlings (Suppl. Fig. 3) showed a delay in the formation of xanthophores, whereas xanthophores were completely and persistently lost in *pax7a; pax7b* double mutant (Suppl. Figs. 5, 6). The melanophore number was not significantly altered in the absence of Pax3 whereas increased in the absence of Pax7 in both medaka and zebrafish (Fig. 2 G, Supp. Fig. 5 J). The iridophore phenotype is faint and ambiguous in both medaka and zebrafish (Suppl. Figs. 4 B, D, F, 6 C, F, I, K)

The mutation in the paralogous gene, *pax3a* in medaka and *pax3b* in zebrafish, did not affect pigment cell development, nor even enhance the phenotypes due to the single paralogous mutation (medaka *pax3b^-/-^*and zebrafish *pax3a^-/-^*) (Suppl. Figs. 7 and 8).

In summary, the phenotypes due to loss of Pax3, by *pax3b* in medaka and *pax3a* in zebrafish, are most evident in xanthophores/leucophores and not significant or less severe in other cell types; notably they do not include obvious melanophore phenotypes. Our results suggest that Pax3 is involved in the specification of xanthophores and leucophores in medaka, and xanthophores in zebrafish (which lack leucophores).

### Loss of Pax3 affects the expression of early xanthophore/leucophore markers

To investigate the early phenotypes of pigment cells in the absence of Pax3, we examined the expression pattern of specification markers. We used *GTP cyclohydrolase I (gch)* as a marker for xanthophore and leucophore progenitors (Pelletier et al., 2001), *dct* as a marker for melanophore progenitors (Kelsh et al., 2000) and *purine nucleoside phosphorylase (pnp4a)* as a marker for iridophore progenitors (Kimura et al., 2017; Petratou et al., 2021; Petratou et al., 2018).

In medaka, expression of *gch* in xanthophore/leucophore progenitors were severely lost in the *pax3b* mutant and completely absent in the *pax7a* mutant (Fig. 3 A, B, C). Melanophore progenitors expressing *dct* appeared to be slightly reduced in number in the *pax3b* mutant whereas they are unaltered or even increased in number and expression level compared to WT in the *pax7a* mutant (Fig. 3 D, E, F). Iridophore progenitors expressing *pnp4a* appeared to be reduced in the yolk sac but not in the eyes in the *pax3b* mutant, whereas they are unaltered in the *pax7a* mutant (Fig. 3 G, H, I). We obtained roughly similar results from zebrafish *pax3a* and *pax7a; pax7b* mutants (Suppl. Fig. 9), except that *dct*-expressing melanophore progenitors are little changed by the loss of Pax3 (Suppl. Fig. 9 D, E) and *pnp4a*-expressing iridophore progenitors appear to be slightly increased in the *pax7a; pax7b* mutant (Suppl. Fig. 9 G-I).

**Fig. 3.**
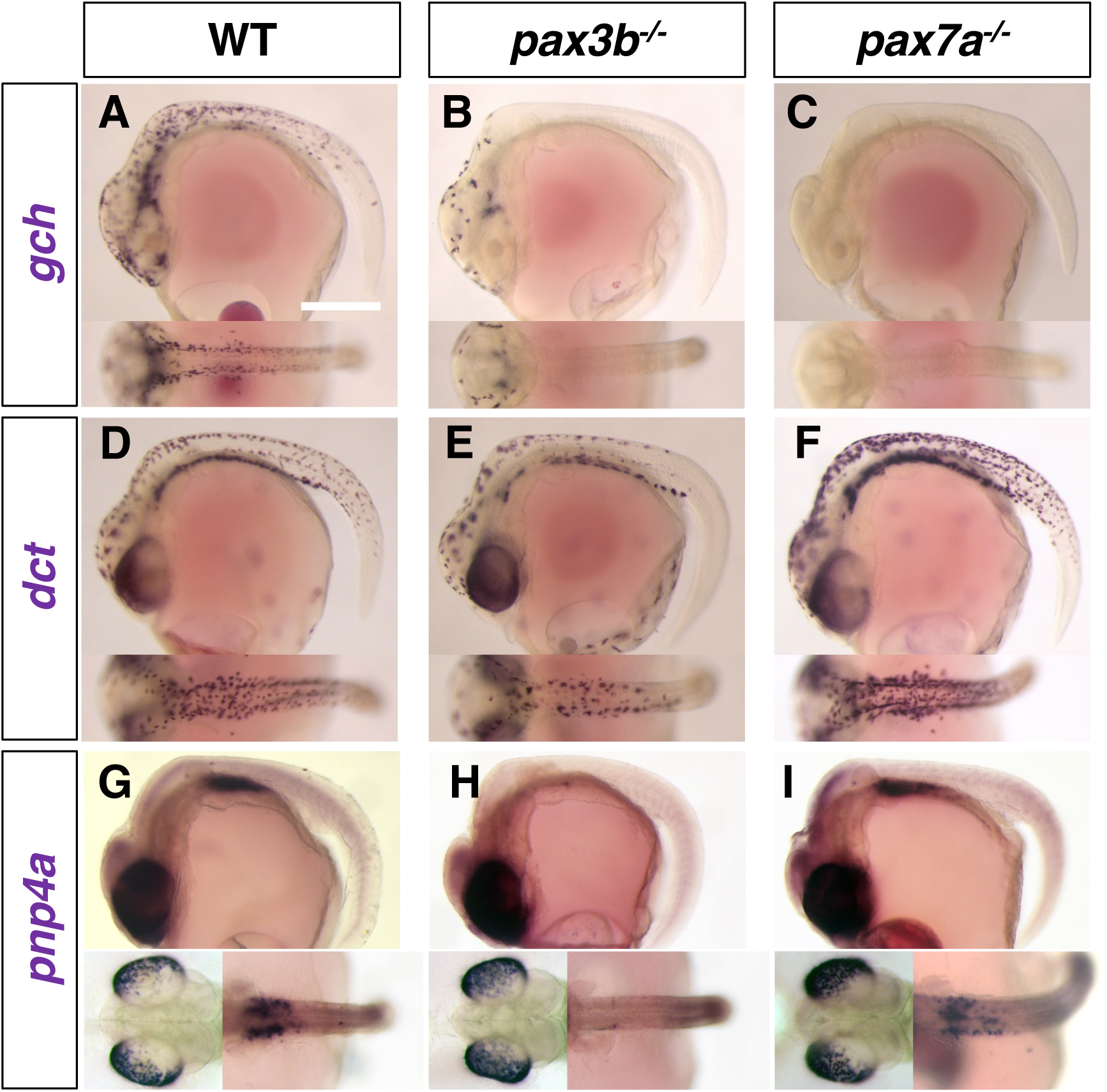
Pigment cell progenitors in medaka *pax3b* and *pax7a* mutant embryos. (A, B, C) *gch*. (D, E, F) *dct*. (G, H, I) *pnp4a*. (A, D, G) WT. (B, E, H) *pax3b^-/-^* mutant. (C, F, I) *pax7a^-/-^* mutant. (A-L) Lateral views at top and dorsal views at bottom at stage 29. Scale bar = 250 µm The *gch*-expressing progenitors of xanthophore and leucophore are severely decreased in *pax3b* mutant (A, B), with only a few remaining anteriorly, and completely absent in *pax7a* mutant (C), which is consistent with the severity of the phenotypes in these mutants at later (hatching) stages. The *dct-*expressing progenitors of melanophore are slightly decreased in number in *pax3b* mutant (D, E) while unaltered or even increased in number and expression level in *pax7a* mutant compared to WT (F). The *pnp4a*-expressing progenitors of iridophore are not altered in the eyes in both mutants (G, H, I, bottom left), but the putative progenitors of peritoneum iridophores are decreased in *pax3b* mutant (G, H) but not in *pax7a* mutant (I, bottom right).

Like the phenotypes of pigmented cells in hatchlings, those of early specification markers in embryos suggest that the defects due to loss of Pax3 (Pax3a in zebrafish and Pax3b in medaka) are most manifest in the formation of xanthophores/leucophores.

### *pax3* functions upstream of *pax7*

To investigate the genetic relationship between *pax3b* and *pax7a*, we examined the expression patterns of both *pax3b* and *pax7a* in the *pax7a* and *pax3b* mutants. In situ analyses showed that while *pax3b*-expressing cells were unaffected in the *pax7a* mutant (Fig. 4A, B), *pax7a*-expressing NCCs were absent in *pax3b* mutant embryos (Fig. 4C, D). Similarly, in zebrafish, both *pax7a*– and the *pax7b*-expressing cells were lost in the *pax3a* mutant (Fig. 4 E-J). The results suggest that, consistent with their expression timings, in the NCCs *pax3b* functions upstream of *pax7a* in medaka, and that *pax3a* functions upstream of *pax7a* and *pax7b* in zebrafish.

**Fig. 4.**
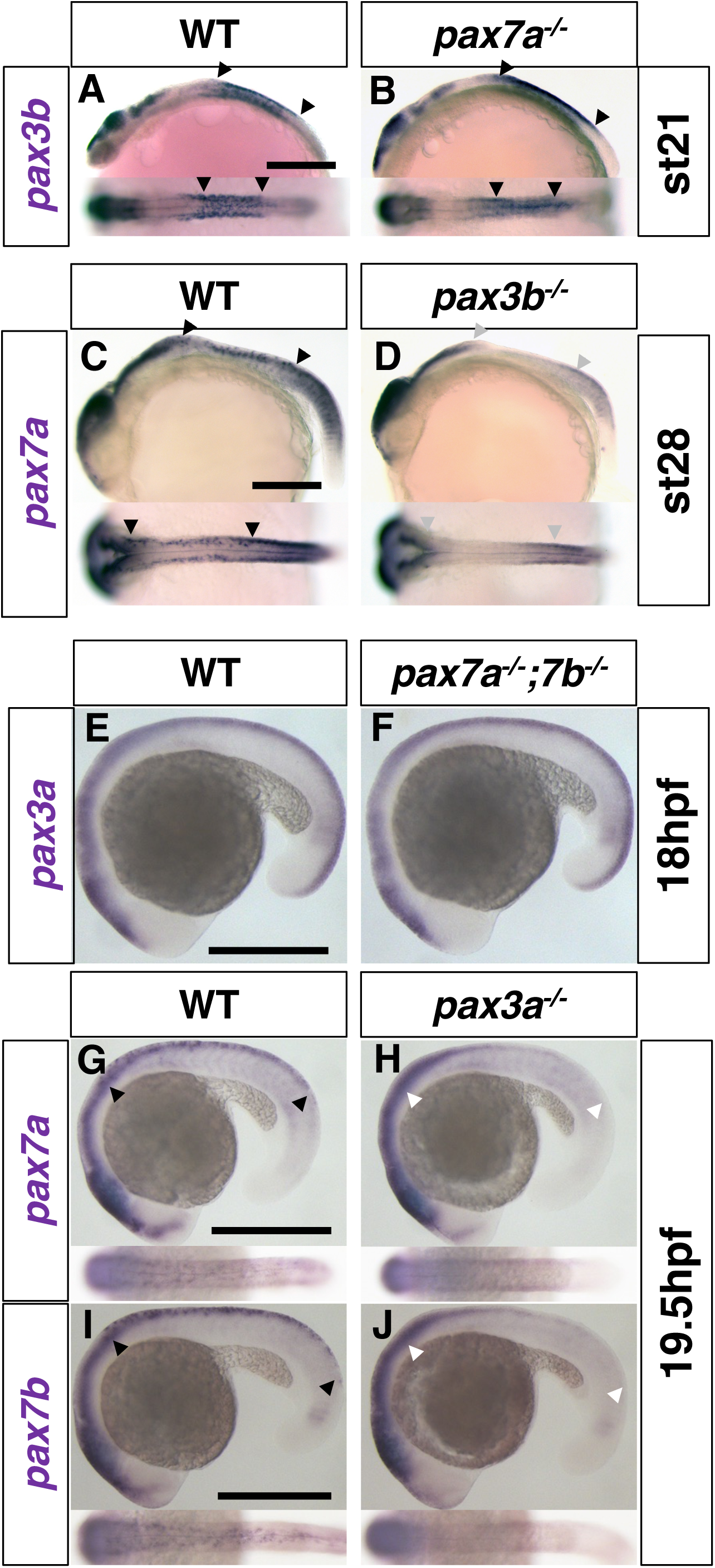
p*a*x3 functions upstream of *pax7*. (A-D) Medaka. (A, B) *pax3b* expression at stage 21 (6-somite). (C, D) *pax7a* expression at stage 27.5 (27-somite). (E-J) Zebrafish. (E, F) *pax3a* expression at 18 hpf (18-somite). (G, H) *pax7a* expression at 19.5 hpf (21-somite). (I, J) *pax7b* expression at 19.5 hpf (21– somite). In medaka, *pax3b* expression is similar in WT and *pax7a* mutant embryos (A, B) whereas *pax7a* expression is largely lost from NCCs in *pax3b* mutant embryo compared to WT (C, D). Similarly, in zebrafish, *pax3a* expression is not altered in *pax7a; pax7b* double mutant embryo (E, F), but *pax7a* and *pax7b* expression is largely lost from NCCs in *pax3a* mutant embryo compared to WT (G-J). Lateral views at top and dorsal views at bottom. Arrowheads indicate the anterior and posterior ends of the expression in the neural crest. Gray and white arrowheads indicate the absence of expression. Scale bar = 250 µm

### The expression of *mitf* is dependent on *pax3*

The *dct* expression patterns in zebrafish and medaka Pax3 mutants are intriguing, because they suggest that the core role for Pax3 in melanocyte development in mammals has been conserved in medaka, but perhaps not in zebrafish, melanophores. To test directly the idea that the core GRN consisting of the transcription factors Sox10, Pax3 and Mitf plays a central role in melanophore development in teleosts, we assessed whether *mitf* expression might be dependent on Pax3 in medaka and zebrafish, similar to its role in mammals (Lang et al., 2005), by examining *mitfa* expression in medaka *pax3b* and zebrafish *pax3a* mutant embryos (Fig. 5). The *mitfa*-expressing cells were severely reduced in stage 28 medaka *pax3b* mutants, whereas comparable or rather increased in *pax7a* mutant compared to wild-type embryos (Fig. 5 A, B, C). Similarly in zebrafish at 19.5 hpf, early *mitfa* expression was partially lost in *pax3a* mutant but not in *pax7a; pax7b* double mutant embryos (Fig. 5 G, H, I). The expression pattern in zebrafish in particular indicates that the Pax3 mutation might cause delayed upregulation of *mitfa,* whereas the loss of Pax7 activity might allow precocious upregulation of *mitfa*. Overall, and importantly, these results indicate that loss of Pax3 causes a reduction in *mitfa* expression in medaka and zebrafish, suggesting that *mitfa* transcription is partially dependent on Pax3 activity.

**Fig. 5.**
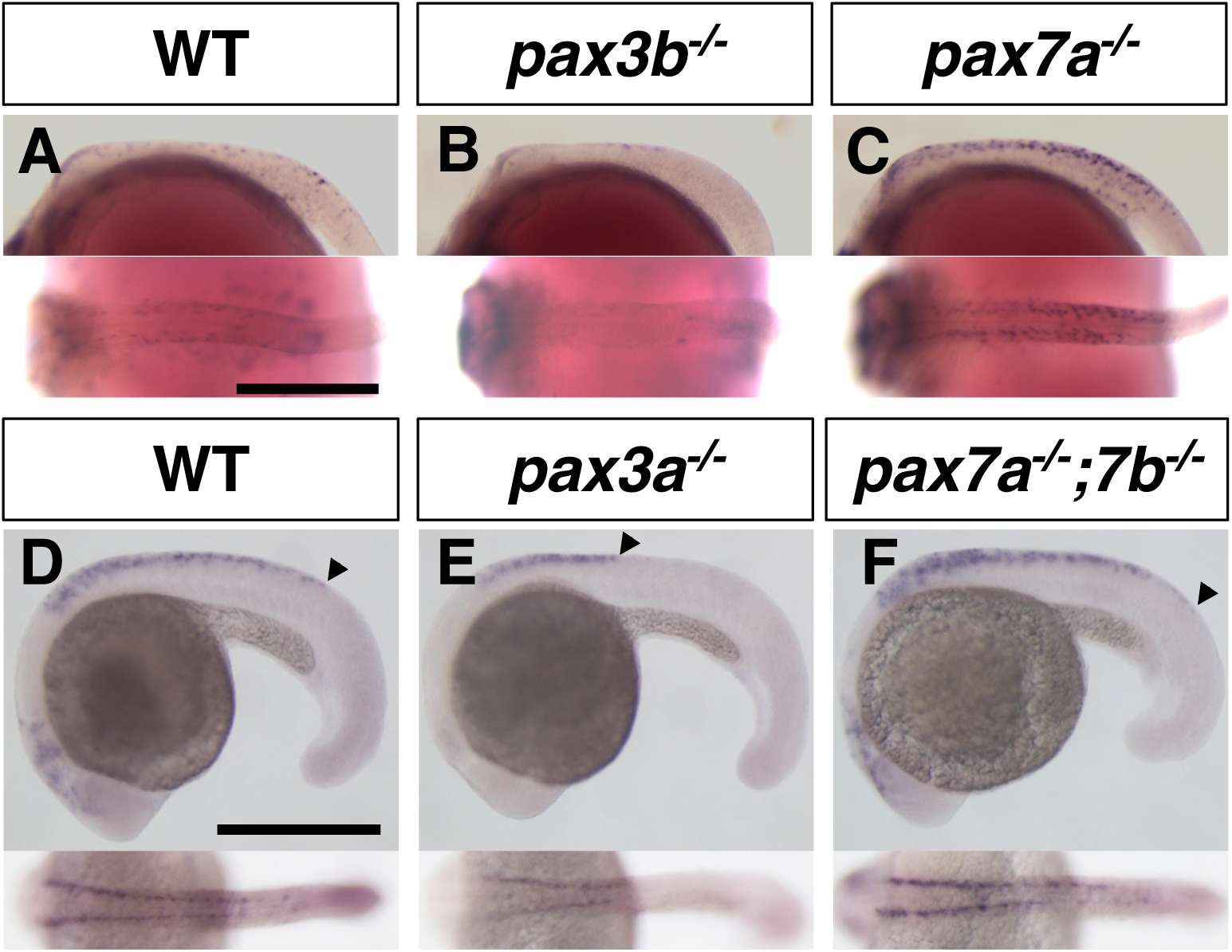
Loss of *pax3* leads to decrease in *mitfa* expression. (A-C) Medaka embryos, stage 28 (30-somite). (A) WT. (B) *pax3b^-/-^* mutant. (C) *pax7a^-/-^*mutant. (D-F) Zebrafish embryos, 19.5 hpf (21-somite). (D) WT. (E) *pax3a^-/-^* mutant. (F) *pax7a^-/-^*; *pax7b^-/-^* mutant. Arrowheads indicate the posterior ends of *mitfa* expression in the neural crest (G-I). Lateral views at top and dorsal views at bottom. Scale bar = 250 µm The *mitfa*-expressing cells are largely lost in the absence of Pax3 in medaka (A, B) and in zebrafish (D, E). Those cells appear to be normal or rather increased in medaka *pax7a* mutant (C) and in zebrafish *pax7a; pax7b* double mutant (F).

### Cooperative function of Pax3 and Sox10 can promote *mitf* expression

To investigate whether Pax3 directly controls *mitf* transcription in teleosts, we performed a Pax3 overexpression experiment by synthetic RNA injection into zebrafish embryos. Working with the assumption that Pax3 cooperates with Sox10 to activate *mitf* transcription, as in mice, we injected synthetic RNAs of *pax3a* and/or *sox10* into one-cell stage embryos and examined whether *mitfa* mRNA was ectopically expressed in the injected embryos at 6 hours post fertilization (hpf), when *mitfa* mRNA is not yet endogenously expressed in zebrafish (Fig. 6 A, A’).

**Fig. 6.**
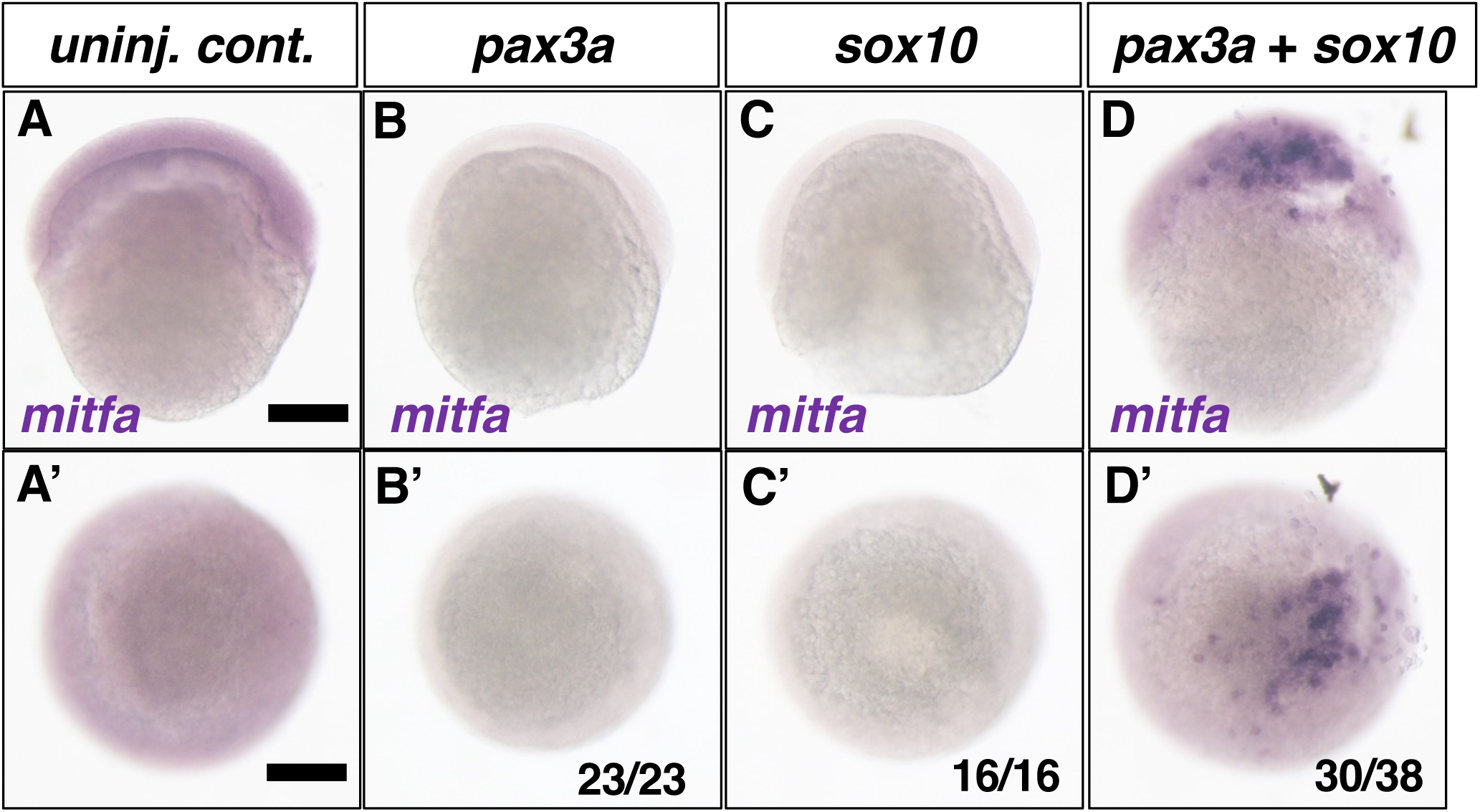
Overexpression of *pax3a* and *sox10* induces ectopic expression of *mitfa* in zebrafish. (A-D, A’-D’) *mitfa* expression. 6 hpf. (A, A) Control embryo. (B, B’) *pax3a* synthetic RNA injected embryo. (C, C’) *sox10* synthetic RNA injected embryo. (D, D’) *pax3a* and *sox10* synthetic RNA co-injected embryo. (A-D) Lateral views. (A’-D’) Animal pole views. Representative embryos are shown. Fractions indicate the number of embryos with the representative phenotype shown over the total injected. *mitfa* mRNA is not expressed endogenously in 6 hpf embryo (A). While overexpression of *pax3a* or *sox10* by synthetic RNA injection into 1– to 2-cell stage embryo fails to induce *mitfa* expression at 6 hpf, simultaneous overexpression of *pax3a* and *sox10* can induce ectopic expression of *mitfa*. Scale bar = 200 µm

While *mitfa* mRNA was not detected in situ when Pax3a or Sox10 were overexpressed alone (Fig. 6 B, B’, C, C’, 23/23 and 16/16), simultaneous overexpression of these transcription factors ectopically induced *mitfa* expression at a high rate (Fig. 6 D, D’, 30/38). Our results suggest that the cooperative action of Pax3 and Sox10 to promote *mitf/Mitf* expression is conserved between zebrafish and mice (Lang et al., 2005).

### Mitfa can drive the progenitor markers for melanophore and xanthophore

Our results above suggest that the GRN consisting of Sox10, Pax3 and Mitf is conserved in zebrafish, where Mitfa plays an essential role in melanophore differentiation (Greenhill et al., 2011). On the other hand, loss of Pax3 has the most severe effect on xanthophore formation in zebrafish and medaka. Given that Mitf mediates Pax3 function, we hypothesized that Mitf is involved in regulating not only melanophore differentiation but also xanthophore differentiation in zebrafish. To test this, we examined whether *mitfa* overexpression could induce ectopic expression of markers for melanophore (*dct*) and xanthophore (*gch*) progenitors in zebrafish embryos (Fig. 7). The *dct* and *gch* mRNAs were not expressed endogenously in control 6 hpf embryos (Fig. 7 A, D). Injection of *mitfa* synthetic RNA induced ectopic mRNA expression of *dct* (Fig. 7 B, 22/23), but also of *gch*, (Fig. 7 E, 20/24). Thus, it is likely that Mitfa can promote both xanthophore and melanophore fates in zebrafish when overexpressed.

**Fig. 7.**
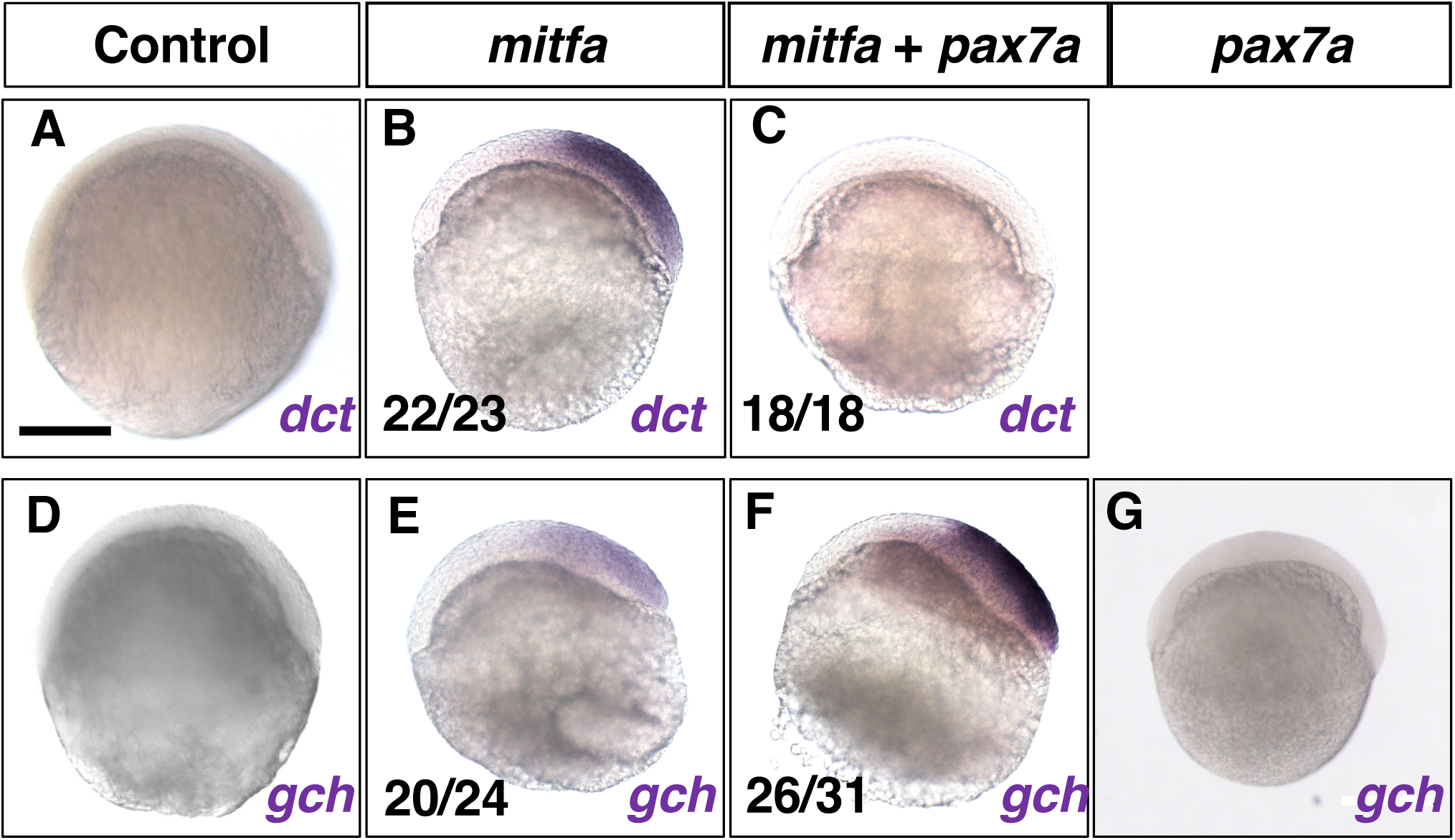
Overexpression of *mitfa* induces ectopic expression of progenitor markers for melanophore and xanthophore. (A-C) *dct* expression. (D-G) *gch* expression. 6 hpf. (A, D) Control embryos. (B, E) *mitfa* synthetic RNA injected embryos. (C, F) *mitfa* and *pax7a* synthetic RNA injected embryos. (G) *pax7a* synthetic RNA injected embryo. Overexpression of *mitfa* alone can strongly induce ectopic *dct* expression (A, B). Simultaneous overexpression of *pax7a* suppresses the ectopic *dct* expression by *mitfa* (C). Ectopic expression of *gch* mRNA is induced by overexpression of *mitfa* alone (D, E) and more strongly induced by overexpression of *mitfa* and *pax7a* in combination (F). Overexpression of *pax7a* alone is unable to induce ectopic *gch* expression (G).

### Pax7 can suppress Mitf action to drive *gch* transcription

What determines the fate choice of the *mitfa*-expressing progenitors to either melanophore or xanthophore fate? To address this, we focused on the previous report in mice that Pax3 can inhibit Mitf-mediated transcriptional activation of *Dct* after activating *Mitf* transcription in cooperation with Sox10 (Lang et al., 2005). We hypothesized that after Pax3 has activated *mitfa* expression, Pax7 instead of Pax3 might play the inhibitory role against Mitfa-driven transcription of *dct* in zebrafish. To test this hypothesis, we examined whether Pax7 could affect the ectopic expression of *dct* and *gch*, when co-overexpressed with Mitfa in zebrafish embryos. The ectopic expression of *dct*, which was detected in the *mitfa*-injected embryos, was lost in the 6 hpf embryos co-injected with *mitfa* and *pax7a* RNAs (Fig. 7 C, 18/18). By contrast, *pax7a* co-injection enhanced the ectopic expression of *gch* in the *mitfa*-injected 6 hpf embryos (Fig. 7 F, 26/31). Interestingly, injection of *pax7a* synthetic RNA alone was unable to activate ectopic *gch* expression (Fig. 7 G).

### Mitf specifies the fate of melanophore and xanthophore in medaka

To determine if Mitf activity is required not only for melanophore fate but also for xanthophore fate, we attempted to generate loss-of-function mutants for Mitf in medaka. We found two *mitf* paralogous genes, *mitfa* and *mitfb*, in the medaka genome as in zebrafish (Lister et al., 2001). The expression pattern of these genes, examined by in situ, showed that both appeared to be expressed in the NCCs (Suppl. Fig. 10). We therefore induced mutations in *mitfa* and *mitfb* using CRISPR/Cas9, targeting the region encoding the basic helix-loop-helix domain (Suppl. Fig. 11).

We successfully generated mutations in the *mitfa* and *mitfb* genes in medaka (Suppl. Fig. 11). Single loss of *mitfb* resulted in a defect in melanophore formation, being delayed at 2 dpf and partially restored at 3 dpf and at hatching, whereas loss of *mitfa* did not, suggesting that *mitfb* but not *mitfa* plays a crucial role in melanophore development in medaka (Fig. 8 A, B, D, E, G, H, not shown for *mitfa* mutant). The double loss of *mitfa* and *mitfb* resulted in complete absence of not only melanophores but also xanthophores and leucophores throughout embryogenesis, and even later during adulthood (Fig. 8 C, F, I, J-M). Iridophore formation appeared normal throughout life (Fig. 8 I2, I3, L, M). These data suggest that Mitfa and Mitfb are redundant in the NCCs and are essential for the development of melanophores, xanthophores and leucophores in medaka.

**Fig. 8.**
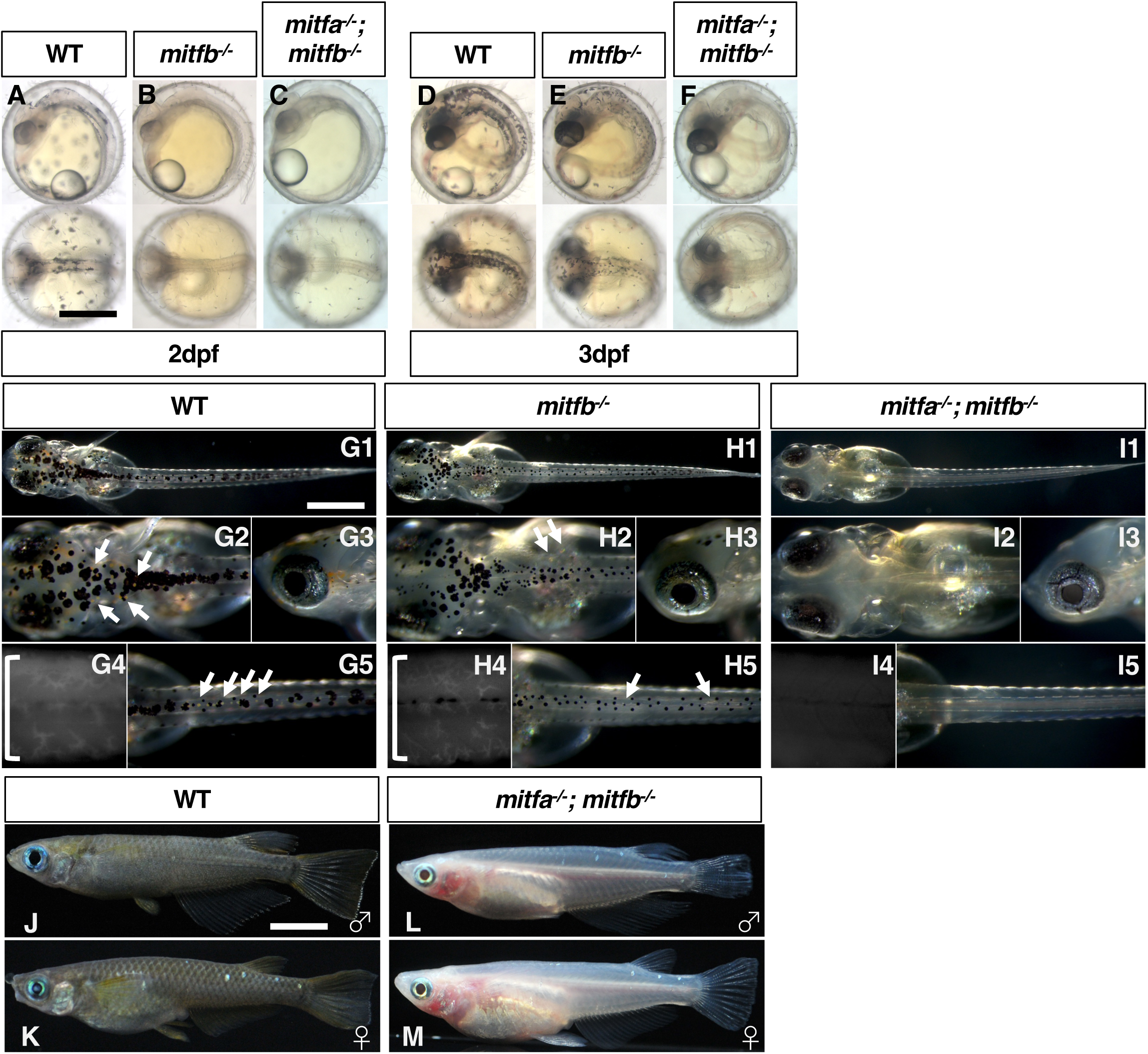
Phenotypes of medaka *mitfa* and *mitfb* double homozygotes. (A-C) 2 dpf embryos. (D-F) 3 dpf embryos. (G-I) 7 dpf hatchlings. (J-M) 4-month-old adult fish. (J, L) males. (K, M) females. (G4, H4, I4) Autofluorescence images showing xanthophores under UV light. Upper panels in A-F and G-I1, 2, 5 are dorsal views. Lower panels in A-F, G-I3, 4 and J-M are lateral views. Melanophores first appear on the head, the anterior body and the yolk at 2 dpf (A) and increase in number to become distributed throughout the body at 3 dpf (D) in the WT embryo. The *mitfa* mutant embryo looks normal at this time (not shown), while the *mitfb* mutant does not have melanophores at 2 dpf (B) but shows their delayed formation at 3 dpf (E). The *mitfa*; *mitfb* double mutant completely lacks melanophores during this period and thereafter (C, F). At the hatching stage, all the four types of pigment cells are differentiated (pigmented) in WT (G1-5). The *mitfb* mutant looks normal except that melanophores are relatively small and few leucophores are found (arrows in G2, G5) compared to those in WT (H1-5). The *mitfa*; *mitfb* double mutant completely lacks not only melanophores (I1, I2, I5) but also xanthophores (I4) and leucophores (I1, I2, I5), but retains iridophores in the eyes and on the yolk (I1, I2, I3). Square brackets indicate xanthophores on the lateral surface of the body (G4, H4). In adulthood, compared to WT (J, K), it is obvious that the *mitfa*; *mitfb* double mutant lacks all visible pigmentation except for that of iridophores in the skin and the iris (L, M). Scale bar = 0.5 mm

In situ analyses support the idea that Mitfs are not only required for melanophore fate but also for specification of xanthophore/leucophore fate (Fig. 9). Expression of *gch* and *dct* is absent from the body surface in the *mitfa*; *mitfb* double homozygotes (Fig. 9 A-D). In contrast, *pnp4a* mRNA is expressed in the eyes and on the yolk similarly in WT and the *mitfa*; *mitfb* double mutant (Fig. 9 E, F). These results are fully consistent with the above phenotype that the double mutant has only iridophores but lacks the other three types of pigment cells.

**Fig. 9.**
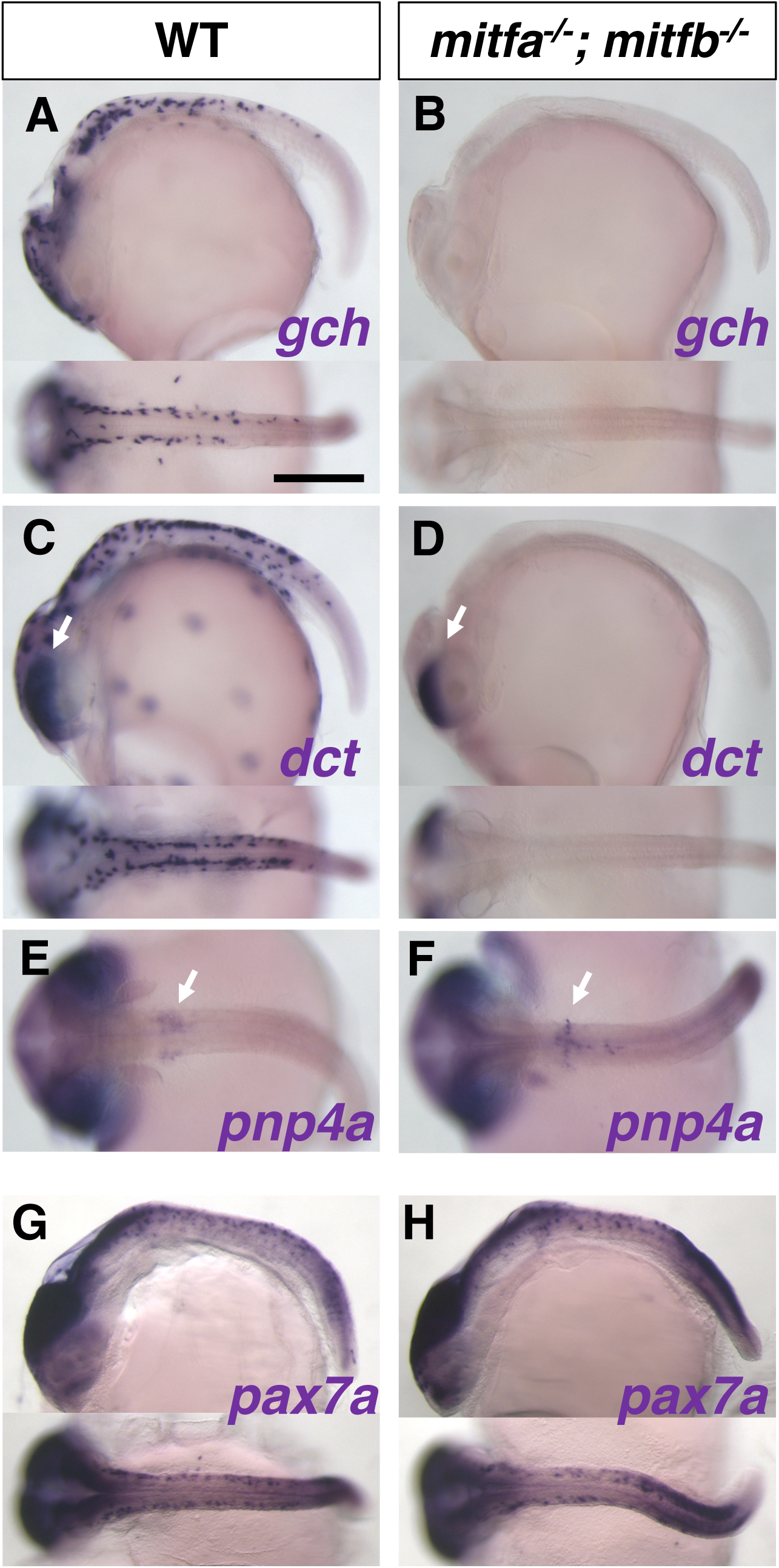
in situ analyses of medaka *mitfa^-/-^*; *mitfb^-/-^* double mutants with pigment cell markers. (A, B) *gch* expression. (C, D) *dct* expression. (E, F) *pnp4a* expression. (G, H) *pax7a* expression. (A, C, E, G) WT. (B, D, F, H) *mitfa^-/-^*; *mitfb^-/-^* mutant. (A-H) Lateral views at top and dorsal views at bottom. (A-D, G, H) Stage 28. (E, F) Stage 29. Scale bar = 250 µm Results of in situ analyses with the specification markers, *dct* for melanophore, *gch* for xanthophore/leucophore, and *pnp4a* for iridophore, are consistent with the phenotypes of differentiated pigment cells. The *gch*-expressing xanthophore/leucophore progenitors are lost in the double mutant (A, B). The *dct*-expressing melanophore progenitors are absent from the body surface, but not from the eyes (retinal pigment epithelium, RPE) in the double mutant embryo (C, D, arrows indicate the signal in RPE), consistent with the phenotype that the mutant retains melanized RPE (see Fig. 8 I1-3). The *pnp4a*-expressing iridophore progenitors appear unchanged in the double mutant compared to WT (E, F, arrows indicate the signal on the yolk). The *mitfa^-/-^*; *mitfb^-/-^* double mutant shows normal expression pattern of *pax7a* mRNA compared to WT (G, H), suggesting that the defect in xanthophore and leucophore formation in the double mutant is not mediated by Pax7a function. Scale bar = 0.5 mm

To exclude the possibility that Mitfs regulate *pax7a* expression to activate the transcription of *dct* and *gch,* we examined the expression of *pax7a* mRNA in the *mitfa*; *mitfb* double homozygotes. The *pax7a*-expressing cells are not notably altered in the double mutant (Fig. 9 H) compared to WT (Fig. 9 G), suggesting that Mitfs directly activate the transcription of *dct* and *gch*, and that, consistent with the overexpression results in Fig. 7, Pax7a alone is not sufficient to drive *gch* transcription.

## Discussion

We established medaka and zebrafish mutants of transcription factors that we expected to be involved in the GRN of pigment cell fate specification. Their phenotypes are summarized in Table 1.

**Table 1.** Transcription factor dependencies of each pigment cell-type 0, no change; +, increase; –, decreased; ––, absent Mel, melanophore; Iri, iridophore; Xan, xanthophore; Leu, leucophre.

| TF | Medaka |  |  |  | Zebrafish |  |  |
| --- | --- | --- | --- | --- | --- | --- | --- |
|  | Mel | Iri | Xan | Leu | Mel | Iri | Xan |
| Pax3a | 0 | 0 | 0 | 0 | 0 | + | delayed |
| Pax3b | 0 | - | delayed | delayed | 0 | 0 | 0 |
| Pax7a/7b | + | 0? | -- | -- | + | 0 | -- |
| Mitfa/b | -- | 0 | -- | -- | -- | + | - |

While Pax7 has been considered a requisite for xanthophore differentiation in medaka and zebrafish and additionally for leucophore differentiation in medaka (Kimura et al., 2014; Minchin and Hughes, 2008; Nord et al., 2016), Pax3, paralogous to Pax7, has not yet been extensively studied in the context of pigment cell development. In this study, we elucidated the function of Pax3 in pigment cell development and proposed that the set of transcription factors Sox10, Pax3 and Mitf regulate the fate of melanophore and xanthophore (plus leucophore in medaka). Furthermore, we found that the choice of melanophore vs. xanthophore/leucophore fate appears to be regulated by the interaction of Mitf and Pax7. In the absence of Pax7, Mitf strongly drives a melanophore progenitor gene *dct,* whereas Mitf and Pax7 cooperatively drive a xanthophore/leucophore progenitor gene *gch*.

### Partially overlapping but sequential function of Pax3 and Pax7

Pax3 is expressed and functions before Pax7 during NCC development, which appears to be temporally restricted to the initial phase of pigment cell progenitor formation. Pax7 expression occurs slightly later than, but overlaps with, Pax3 expression, and is long maintained until a later stage of cell differentiation (Fig 1, Suppl. Fig. 2). Assuming that Pax3 and Pax7 are highly homologous and have similar activities (Pax3a vs. Pax7a/7b= 96.8% similarity; Pax7a vs. Pax7b = 100% in homeodomain of zebrafish proteins), loss of Pax3 might be partially compensated by Pax7 at early stages but loss of Pax7 cannot be compensated by Pax3 at later stages because Pax3 is no longer expressed. The mutant phenotypes support this idea: xanthophores and leucophores are partially formed in the absence of Pax3, whereas these pigment cell types are completely lost in the absence of Pax7, in medaka (Fig. 2, Suppl. Fig. 4). This is also the case for xanthophores in zebrafish (Suppl. Figs. 5, 6).

### The gene regulatory network of Sox10, Pax3 and Mitf in teleosts

Pax3 is required to activate *mitf* expression as shown in medaka *pax3b* mutant and zebrafish *pax3a* mutant with delayed *mitfa* expression (Fig. 5). Overexpression of Pax3 together with Sox10 resulted in ectopic expression of *mitfa* in zebrafish embryos (Fig. 6). As the previous report using human cultured cells showed that Pax3 functions with Sox10 to activate *Mitf* expression (Lang et al., 2005), we propose that a conserved role of Pax3 in vertebrates is to form a core melanophore GRN with Sox10 to regulate Mitf in the NCCs.

The medaka and zebrafish *pax3* mutants only showed ambiguously the melanophore– related phenotypes. Although loss of Pax3 appeared to substantially decrease *mitf* expression, it eventually resulted in only partially or barely reduced melanophore formation in both medaka and zebrafish (Fig. 2, Suppl. Figs. 4, 5, 6). We reason this can be explained as *mitf* expression is delayed in the absence of Pax3 and perhaps subsequently compensated by some other transcription factors. In contrast, loss of Pax7 resulted in significant increase of *mitfa*-expression in progenitors and pigmented melanophores in both species (Fig. 2, Suppl. Figs. 4, 5, 6). As Pax7 and presumably Pax3 inhibit Mitf protein from promoting melanophore fate (see below), loss of Pax3 or Pax7 would tend to increase melanophore formation (Fig. 2, Suppl. Fig. 5). These conflicting actions may explain the ambiguous melanophore phenotypes in *pax3* mutants.

### Pax7 interacts with Mitf to segregate pigment cell lineage

In mice, Mitf directly activates *Dct* expression and drives the cells to differentiate into melanocytes (Lang et al., 2005; Ludwig et al., 2004). In teleosts, Mitf also drives *dct* expression, reflecting a broader role in driving melanophore differentiation (Elworthy et al., 2003; Greenhill et al., 2011; Lister et al., 1999). However, here we show that co–expression of Pax7 can inhibit Mitf from driving *dct* transcription, which we suggest has physiological relevance when *pax7* is expressed in a subset of the *mitfa*-expressing neural crest cell population (Fig. 7). In these cells, *gch* is activated by the cooperative action of Pax7 and Mitfa instead of *dct* (Fig. 7), presumably reflecting a broader role for the combinatorial impact of these transcription factors, which together drive differentiation into xanthophores or leucophores. If *pax7* is not co-expressed, the *mitfa*-expressing cells would differentiate into melanophores, as in mice. We still do not know how the *mitfa*– expressing progenitor population is divided into two, one expressing Pax7 and the other not.

## Model

We propose that a more complex GRN than in mammals controls the fate specification of melanophore progenitors and xanthophore/leucophore progenitors in medaka (Fig. 10). Pax3 and Sox10 activate *mitf* expression in the NCC-derived multipotent progenitors for pigment cell. The fate of the *mitf*-expressing progenitor cells is determined by whether or not they express Pax7. These progenitors, if expressing Pax7, give rise to xanthophores and leucophores in medaka (to xanthophores in zebrafish), whereas, if not expressing Pax7, they give rise to melanophores.

**Fig. 10.**
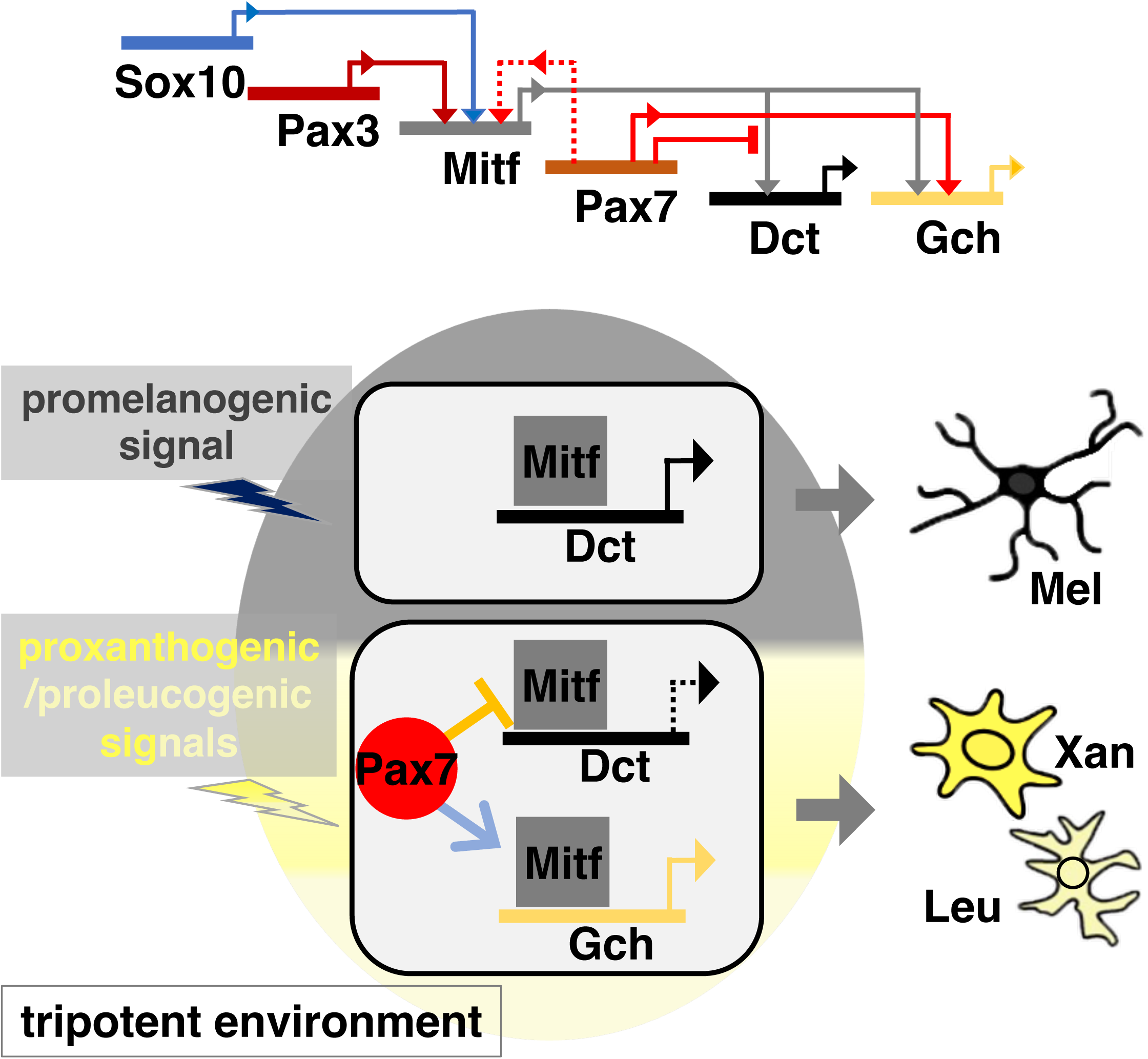
Model. Our results postulate a gene regulatory network that controls the development of melanophores with Dct expression and xanthophores/leucophores with Gch expression. In medaka NCCs, Sox10 (Sox10a and Sox10b in medaka) and Pax3 (Pax3b in medaka) activate Mitf (Mitfa and possibly Mitfb) expression. Mitf can drive transcription of Dct and Gch. Pax7 represses transcription of Dct by inhibiting Mitf, and promotes transcription of Gch cooperatively with Mitf. Pax7 may compensate for loss of Pax3 to activate Mitf expression (dotted line). Expression of Mitf and Pax7 are dependent upon promelanogenic and proxanthogenic/proleucogenic signals respectively (the cell resides in a tripotent environment). A fraction of the Mitf-expressing cells, if expressing Pax7a (lower cell), would differentiate into Gch-expressing xanthophore/leucophore lineage because Mitf and Pax7 cooperatively activate Gch expression while Pax7 represses Dct by inhibiting Mitf. Another cell, not expressing Pax7 (upper cell), would differentiate into Dct-expressing melanophore lineage where Mitf activates Dct expression.

Previously, we and others have explained pigment cell development from NCCs in terms of a progressive fate restriction model, postulating partially-restricted intermediates such as a bipotent progenitor of melanophore and iridophore and a bipotent progenitor of xanthophore and leucophore. Recently, based upon detailed evaluation of pigment cell development in the zebrafish, we have proposed an alternative view, the cyclical fate restriction (CFR) model. Under CFR, we predict that pigment cell development proceeds directly and dynamically from a highly multipotent state (Kelsh et al., 2021; Subkhankulova et al., 2023). We consider that this view is readily applied to the consideration of the situation in medaka too. The key is to replace the concept of a bipotent or tripotent progenitor with that of a bipotent or tripotent environment – one in which the broadly multipotent cell is exposed to two or three key fate-determining signals, which limit the options available to the multipotent cell in that region. In this context, we suggest that a pigment cell progenitor expresses *mitf* in response to a promelanogic environment. But in addition, if the cell receives unknown signals that are xanthogenic and leucogenic (a tripotent environment), and where these are strong/prolonged enough, *pax7* expression becomes dominant, and the cell adopts a xanthophore or leucophore fate. If the signals are too transient/weak, the cell would differentiate into a melanophore.

To our knowledge, Pax7 has not been implicated in pigment cell development in mammals and birds, which lack xanthophores, whereas Pax3 plays a key role in melanocyte development in mammals (Conway et al., 1997; Tassabehji et al., 1992). It is conceivable that the sequential activity of earlier Pax3 and later Pax7 is key to the development of xanthophores and leucophores in vertebrates, and also that loss of Pax3– dependent Pax7 expression may contribute to the evolutionary loss of xanthophore fate from the endothermic vertebrates.

### Mitf is a key player for the formation of both melanophore and xanthophore progenitors

Overexpression of Mitfa alone can ectopically activate the expression of not only *dct* but also *gch* in zebrafish embryos. This suggests an unexpected role of Mitfa in promoting the xanthophore fate. Indeed, our genetic study in medaka showed that Mitfs (Mitfa and Mitfb) are essential for the formation of not only melanophores but also xanthophores and leucophores (Figs. 8, 9).

Mitf has long been believed to be the master regulator of melanophores in teleosts like in mammals (Arnheiter, 2010) since zebrafish *mitfa* mutants lack melanophores completely (Lister et al., 1999). At the same time, however, xanthophore pigmentation becomes reduced in the *mitfa* mutant, implying that Mitf is involved in xanthophore formation and thus that Mitfb compensates for the loss of Mitfa. Lister et al. generated *mitfa; mitfb* double mutants, but did not describe whether they lacked xanthophores (Lister et al., 2001).

Our study clearly shows that loss of Mitfa and Mitfb results in complete absence of melanophores, xanthophores and leucophores throughout life in medaka (Fig. 8), suggesting that Mitfs (Mitfa and Mitfb) are required for fate specification of not only melanophore but also xanthophore and leucophore in medaka. In contrast, zebrafish *mitfa*; *mitfb* double mutants retain xanthophores, as we confirmed by using a novel *mitfb* allele we generated and the *mitfa^nacre^*allele (Suppl. Figs. 12, 13). It remains unclear why Mitf loss-of-function phenotypes differ between zebrafish and medaka with respect to xanthophore formation. However, the slightly enhanced reduction of xanthophores by loss of Mitfb in zebrafish *mitfa/nacre* mutant suggests that Mitf activity is also involved in xanthophore fate development in zebrafish (Suppl. Fig. 13). We speculate that Tfec, a transcription factor homologous to Mitf, and that is required for iridophore specification, might partially compensate for the loss of Mitfa and Mitfb to form xanthophores on the head in zebrafish. Indeed, Tfec and Mitfs are partially redundant for pigment cell development in zebrafish (Petratou et al., 2021; Subkhankulova et al., 2023). Redundancy of Mitf and Tfec may differ between medaka and zebrafish, and the formation of xanthophores in these species may be differentially regulated by Mitf and Tfec.

In conclusion, the core GRN consisting of Sox10, Pax3 and Mitf to regulate pigment cell development is conserved across teleosts and mammals. We propose that the core GRN governs melanophore and xanthophore/leucophore potentials in teleosts. Recruitment of Pax7 to the GRN allows teleosts to create cell diversity, e.g. xanthophore/leucophore, implying an evolutionary aspect of cell diversification in vertebrates. Finally, although Mitf is considered to be a master regulator of the melanocyte phenotype, this picture may be rather misleading outside of a mammalian context; instead we note that our and previous studies make clear that pigment cell fate determination in fish likely reflects the *combinatorial* activity of multiple transcription factors, including those of the Mitf/Tfec, Pax and Sox families. Further work will be required to define fully the transcription factor complements responsible for each cell-type.

## Materials and methods

### Ethics

The animal work in this study was approved by the Nagoya University Animal Experiment Committee and was conducted in accordance with the Regulations on Animal Experiments at Nagoya University.

#### Strains and fish husbandry

The Nagoya and d-rR strains of the medaka fish *Oryzias latipes* were used as the wild type (WT) (Nagao et al., 2014; Yamamoto, 1953). Medaka *pax3a* and *pax3b* mutant strains were generated by TALEN in the d-rR strain and maintained in the d-rR background, designated as d-rR; *pax3a^ex2del5^*and d-rR; *pax3b^ex2del11^*. The d-rR; *pax3b^ex2del11^*strain was crossed with the Nagoya strain to obtain Nagoya; *pax3b^ex2del11^*having melanized melanophores (see Suppl. Figs. 4, 7). The *pax3a^ex2del5/ex2del5^*and *pax3b^ex2del11/ex2del11^* mutants are considered null mutants and are therefore referred to as *pax3a^-/-^* and *pax3b^-/-^*, respectively. Medaka *leucophore free-2* strain, previously described (Kimura et al., 2014), was used as *pax7a* null mutant. Medaka *mitfa^ex6del1^* and *mitfb^ex6del7^* mutation was generated by CRISPR/Cas9 on the Nagoya background. In this paper, the homozygous mutants are designated as *mitfa^-/-^* and *mitfb^-/-^*. The AB strain of the zebrafish *Danio rerio* was used as the wild type (WT). Zebrafish *pax3a, pax3b* and *pax7a* mutant strains were generated by TALEN and *pax7b* mutant was by CRISPR/Cas9. The alleles isolated in this study are *pax3a^ex2del14^*, *pax3b^ex2del11^*, *pax3b ^ex2del16^*, *pax7a^ex2del19^* and *pax7b^ex1del10^*. All the mutant alleles are considered null alleles, and thus these zebrafish mutants are designated as *pax3a^-/-^*, *pax3b^-/-^*, *pax7a^-/-^* and *pax7b^-^/-*.

#### Genotyping

Mutations in medaka and zebrafish were detected using polymerase chain reaction (PCR) fragment length polymorphism by polyacrylamide gel electrophoresis (PAGE), as previously described (Nagao et al., 2018). PCR primer sets are: medaka *pax3a^ex2del5^*, 5ʹ– AGGTCTCTGGATTTTTCTAACCTAAACCCG-3ʹ and 5ʹ-TGGTGCGCCATCTCCACGATCTTATG-3ʹ; medaka *pax3b^ex2del11^*, 5ʹ–TGCTGTGATTGAACGCAGTGTCCACCCCGC-3ʹ and 5ʹ-GTCTCCTGGTACCGGCACAGGATTTTGGAC-3ʹ; zebrafish *pax3a^ex2del14^*, 5ʹ–CGCTGACTTTTCCTCTTTTGT-3’ and 5ʹ-GCGGATGTGATTGGGCAAAGG-3’; zebrafish *pax3b^ex2del11^ ^or^ ^ex2del16^*, 5’-TGTCAACTCCGATGGGTCAG-3’ and 5’–AGAGACTCTGAGCTGGCGGG-3’; zebrafish *pax7a^ex2del19^*, 5’–TCTCAACACCTCTGGGTCAAG-3’ and 5’-CCATTTCCACTATTTTGTGTC-3’; zebrafish *pax7b^ex1del10^*, 5’-GTAGAATGTCATCCTTACCGG-3’ and 5’–CTTCCAAGGGGAATCCGGTGC-3’; medaka *mitfa^ex6del1^*5’–AGGTTCAACATTAATGACCGCATT-3’ (WT allele specific sense) and 5’–AGGTTCAACATTAATGACCGCATA-3’ (Mutant allele specific sense) and 5’–TGATTGCAGATGATTTGCCCATC-3’ (common antisense); medaka *mitfb^ex6del7^*, 5’–GCATTGAAGCATTTCTTCCAG-3’ and 5’-AAAATGGCGGGATATACGTAC-3’. PCR products were separated by PAGE to detect the deletion in size or by agarose gel to check presence or absence (for *mitfa^ex6del1^*).

#### TALEN– and CRISPR/Cas9-mediated mutagenesis

Mutagenesis was done according to the methods described previously, e.g., for TALEN (Ansai et al., 2013) and for CRISPR/Cas9 (Ansai and Kinoshita, 2014). The internal spacer sequence of a pair of TALEN targets are: medaka *pax3a*, 5ʹ– CAACCAGCTCGGCGG-3ʹ; medaka *pax3b*, 5ʹ-TTGCCTAACCATATCCG-3ʹ; zebrafish *pax3a*, 5ʹ-GGGCCGCGTGAACCA-3’; zebrafish *pax3b*, 5’– TCTGCCCAACCACATTC-3’; zebrafish *pax7a*, 5’-TTTTCATCAACGGA-3’.

The target sequences of CRISPR/Cas9 are: zebrafish *pax3b*, 5’– ATCACGGAATCCGGCCCT-3’; zebrafish *pax7b*, 5’-TACCGCGAATGATGCGAC-3’.

We purchased crRNA for targeting medaka *mitfa* and *mitfb*: *mitfa*, 5’– AGGTTCCTAGTTCCTTAATGCGG-3’; *mitfb*, 5’-GAAGGTTCAACATCAACGATCGG-3’ (Integrated DNA Technologies, Iowa, USA), and injected each with tracrRNA and Cas9 protein (Integrated DNA Technologies, Iowa, USA) into one-cell embryos.

#### Transcript detection in whole mount embryos

Whole mount *in situ* hybridization was performed as previously described (Nagao et al., 2014). The digoxigenin (Roche Diagnostics GmbH, Mannheim, Germany)-labeled antisense riboprobe was synthesized from the plasmid containing the full or partial length open reading frame of the cDNAs using SP6 or T7 polymerase (Promega, Madison, WI, USA) after restriction enzyme digestion (New England Bio Labs, Beverly MA, USA).

#### Microscopy

Melanophores were examined under a stereomicroscope (MZ APO, Leica Microsystems, Wetzlar, Germany) using a combination of bright– and darkfield illumination. Leucophores and iridophores were identified in dark field. Xanthophores were identified by detecting their autofluorescence under UV light with a DAPI filter (Imager.D1, Carl Zeiss, Oberkochen, Germany). Zeiss LSM700 confocal microscope was used to observe GFP fluorescent cells in the transgenic embryos.

#### Constructs and microinjection of synthetic RNAs

The full-length open reading frame of the cDNA for each gene was subcloned into pCS2 plasmid (Turner and Weintraub, 1994). Artificial mRNA (synthetic RNA) was transcribed in vitro using SP6 polymerase (Promega, Madison, WI, USA) after NotI digestion (New England Bio Labs, Beverly MA, USA).

## Acknowledgements

The authors thank Dr. Masato Kinoshita for gifts of TALEN and CRISPR/Cas9 tools.

## Figure legends

**Suppl. Fig. 1.**
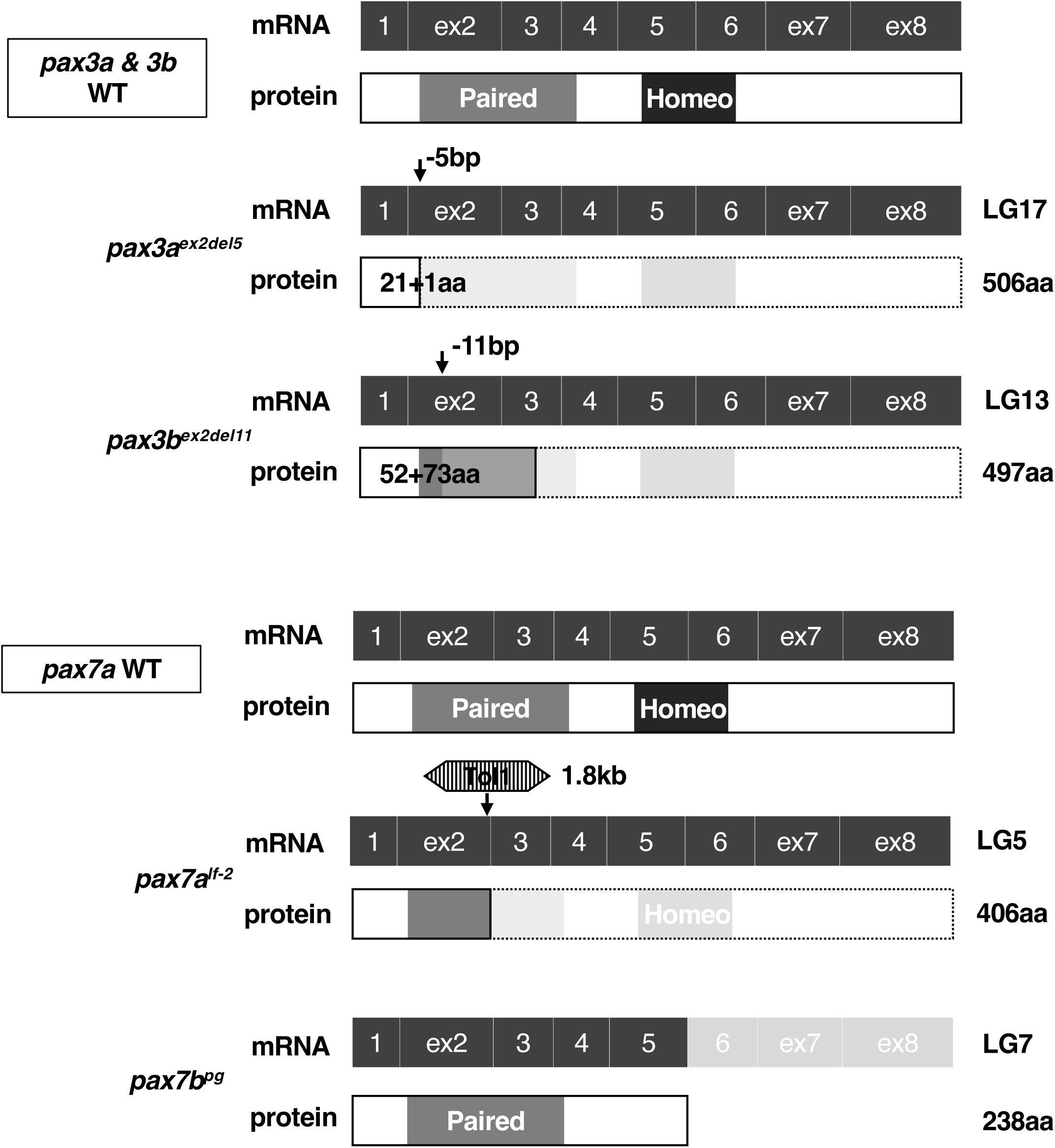
The *pax3* and *pax7* mutations in medaka. We generated mutants for *pax3a*, *pax3b* and *pax7b* in medaka using CRISPR/Cas9 or TALEN. We selected a target in the region encoding the paired domain. A five base pair deletion in the second exon of *pax3a* gene (*pax3a^ex2del5^*) and an eleven base pair deletion in the third exon in *pax3b* gene (*pax3b^ex2del11^*) were generated in medaka. All the mutants were viable and fertile, and so we successfully established each strain. Schematics of predicted primary structures are shown for wild type (WT) gene or mRNA and protein and mutant protein of Pax3a, Pax3b, Pax7a and Pax7b. The linkage group (LG) where the gene is located is shown on the right to the mRNA. The size (aa, amino acid) of the WT protein is shown to the right to the protein. The position and size (bp) of the deletion or insertion mutation is indicated by arrow in the corresponding exon. The size of the mutant protein is given as the endogenous amino acid sequence + a de novo sequence following the frame-shift due to the deletion mutation (e.g. 21 + 1aa means that the Pax3a^ex2del5^ protein retains 21 N-terminal amino acids and one de novo amino acid). The *pax7a* mutation is an insertional mutation of Tol2 transposon (1.8kb) at the end of exon 2, previously known as *leucophore free-2* mutant (Kimura et al., 2014). Pax7b was found in the medaka genome database as a pseudogene lacking exons 6-8 (ENSORLG00000023957). None of the mutant proteins has a complete paired box (gray) nor a homeobox (black).

**Suppl. Fig. 2.**
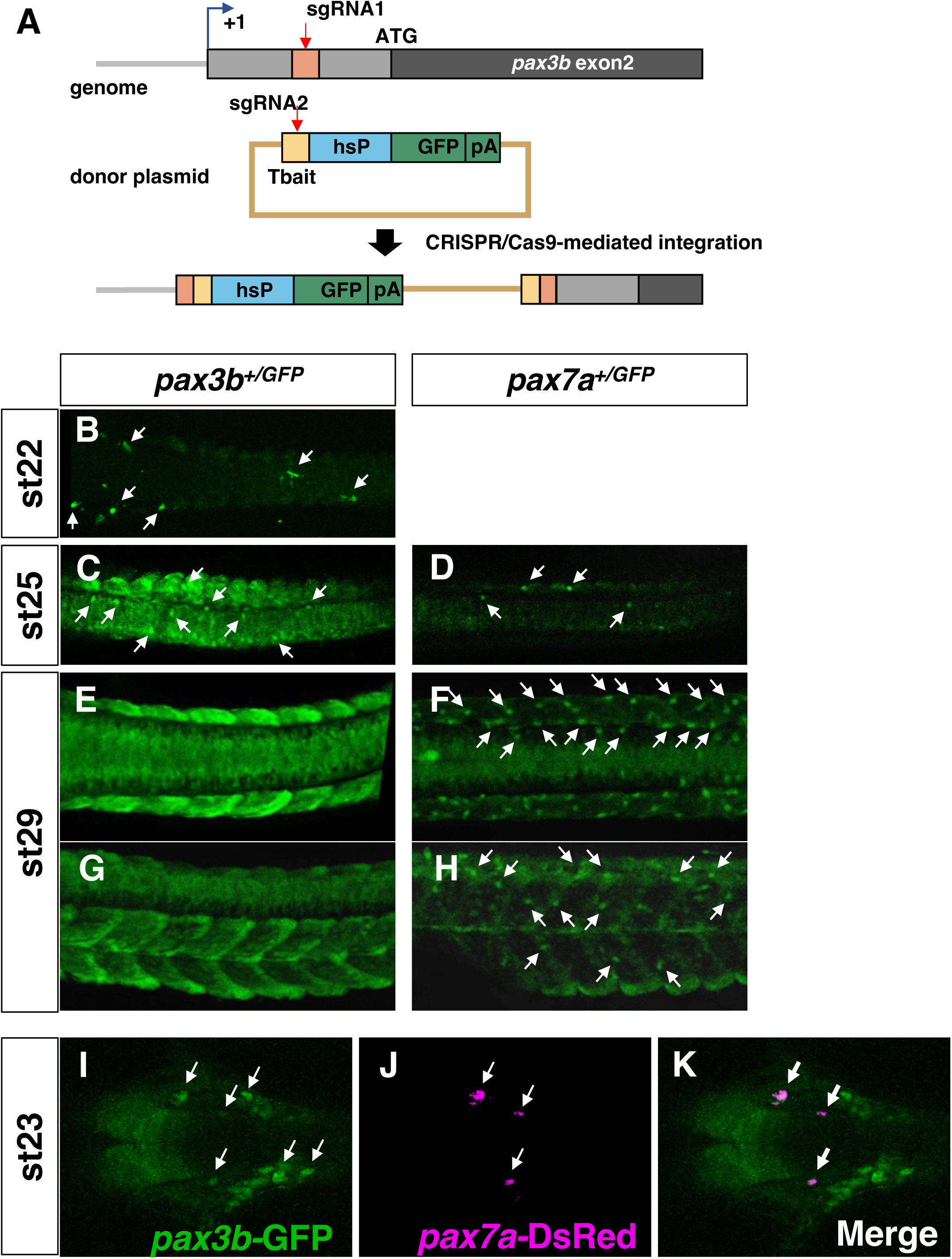
Labeling *pax3b*– and *pax7a*-expressing cells in medaka GFP-knock-in transgenic lines. (A) Method for targeted integration of the GFP cDNA using CRISPR/Cas9. sgRNA1 was designed to target a sequence (5’-GGCTAGACAGCAGTGTCCC-3’) upstream of the *pax3b* start codon. The sgRNA2 was used to cleave of the donor plasmid containing Tbait sequence upstream of the insertion cassette (hsP-GFP-pA) (Watakabe et al., 2018). (B, C, E, G, I, K) *pax3b^+/GFP^* knockin line Tg(*pax3b*-*hs:GFP*). (D, F, H, J) *pax7a^+/GFP^* knockin line Tg(*pax7a*-*hs:GFP*). (J, K) Tg(*pax3b*-*hs:GFP*); Tg*BAC*(*pax7a*-*DsRed*). Tg(*pax3b*-*hs:GFP*) has been established and is reported for the first time in this study. Tg(*pax7a*-*hs:GFP*) and Tg*BAC*(*pax7a*-*DsRed*) transgenic lines have been reported previously (Nagao et al., 2018; Watakabe et al., 2018). The *pax3b*-expressing cells, labeled with GFP fluorescence, are observed in premigratory NCCs, neural tissues and somites at stages 22-25 (B, C). Later than these stages, the fluorescence has disappeared from the NCCs while signals are still observed in the somites and neural tube (E, G). A few cells in premigratory NCCs are positive for fluorescence in the *pax7a^+/GFP^* embryo at stage 25 (D). The *pax7a*-expressing cells are located in migrating NCCs, expanding ventralaterally (F, H). The cells that are double positive for GFP and DsRed in The Tg(*pax3b*-*hs:GFP*); Tg*BAC*(*pax7a*-*DsRed*) embryo at stage 23 (arrows) indicate that those express both Pax3b and Pax7a. At the same time, the cells positive only for GFP are observed (I, J, K).

**Suppl. Fig. 3.**
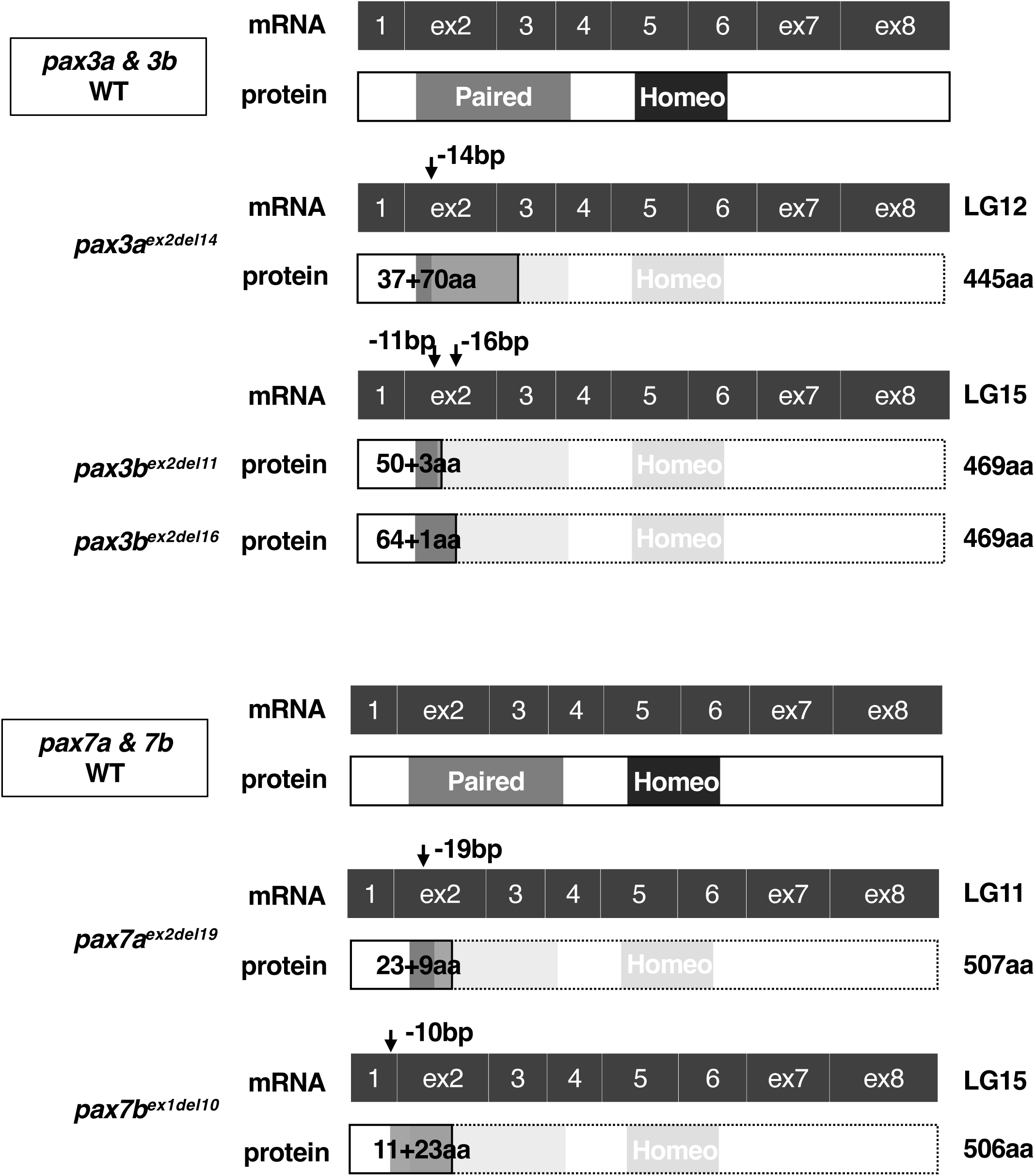
The *pax3* and *pax7* mutations in zebrafish. We generated mutants for *pax3a*, *pax3b*, *pax7a* and *pax7b* in zebrafish using CRISPR/Cas9 or TALEN. We selected a target in the region encoding the paired domain. A fourteen base pair deletion in the second exon of *pax3a* gene (*pax3a^ex2del14^*), an eleven base pair deletion in the second exon of *pax3b* gene (*pax3b^ex2del11^*), a nineteen base pair deletion in the second exon of *pax7a* gene (*pax7a^ex2del19^*) and a ten base pair deletion in the first exon of *pax7b* gene (*pax7b^ex1del10^*) were generated in zebrafish. All the mutants were viable and fertile, and so we successfully established each strain. In addition, the double mutants of the paralogous genes, *pax3a; pax3b* double and *pax7a; pax7b* double, were also viable and fertile, and were used for our analyses when necessary. Schematics of predicted primary structures are shown for wild type (WT) mRNA and protein and mutant protein of Pax3a, Pax3b, Pax7a and Pax7b. The linkage group (LG) where the gene is located is shown to the right of the mRNA. The size (aa, amino acid) of the WT protein is shown to the right of the protein. The position and size (bp) of the deletion mutation is indicated by arrow in the corresponding exon. The size of the mutant protein is given as the endogenous amino acid sequence + a de novo sequence following the frame-shift due to the deletion mutation (e.g., 37 + 70aa means that Pax3a^ex2del14^ protein retains 37 N-terminal amino acids and 70 de novo amino acids). None of the mutant proteins has a complete paired box (gray) nor a homeobox (black).

**Suppl. Fig. 4.**
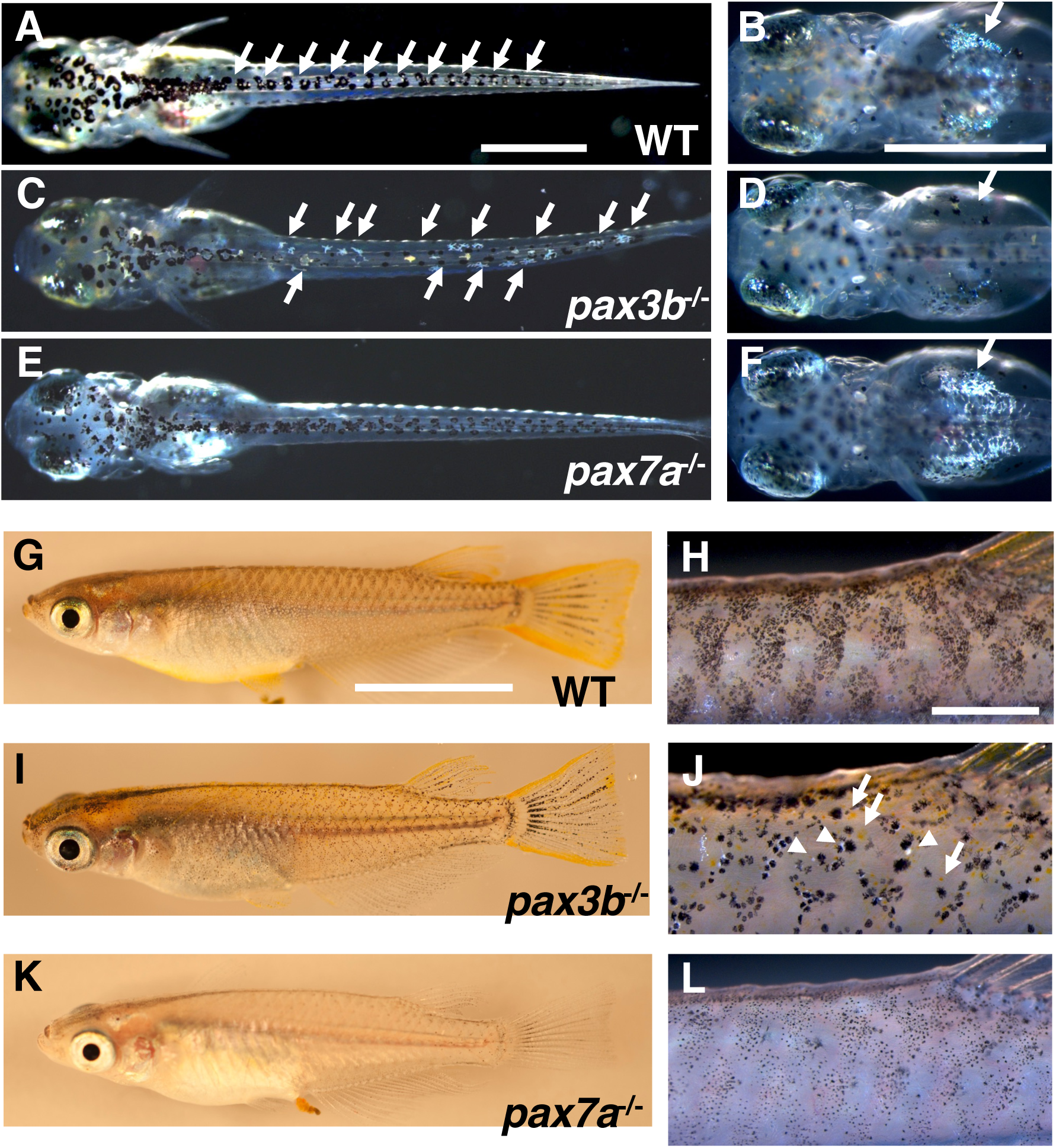
Larval and adult gross phenotypes of medaka *pax3b* and *pax7a* mutants. (A, C, E) 8 dpf hatchlings. (A) Wild type (WT). (C) *pax3b^-/-^*. (E) *pax7a^-/-^*. (B, D, F) Close-up views of the dorsal region of the yolk in 7 dpf hatchlings. (B) WT. (D) *pax3b^-/-^*. (F) *pax7a^-/-^*. (G-L) 4-month-old adult fish. (G, H) WT. (I, J) *pax3b^-/-^*. (K, L) *pax7a^-/-^*. (H, J, L) Close-up views of the anterior region of the dorsal fin of the fish in G, I, and K, respectively. (A-F) Dorsal views in dark field. (G-L) Lateral views. Scale bars = 0.3 mm (A-F), 2 cm (G, I, K) and 0.5 mm (H, J, L) Melanophores, leucophores and iridophores seem only slightly reduced in *pax3b^-/-^*hatchlings (A-D, arrows indicate leucophores and iridophores). See Fig. 2 B, D, F for xanthophores. These pigment cells appeared to have partially recovered by adulthood (arrows indicate xanthophores and arrowheads indicate leucophores in J), although melanophores look fewer and enlarged in the *pax3b* mutant (G-J). In the *pax7a* mutant, leucophores and xanthophores are completely absent at hatching stages (E, see also Fig. 2 F) and during adulthood (K, L). Formation of melanophores and iridophores looks relatively unaffected by loss of Pax7a (E, F, K, L).

**Suppl. Fig. 5.**
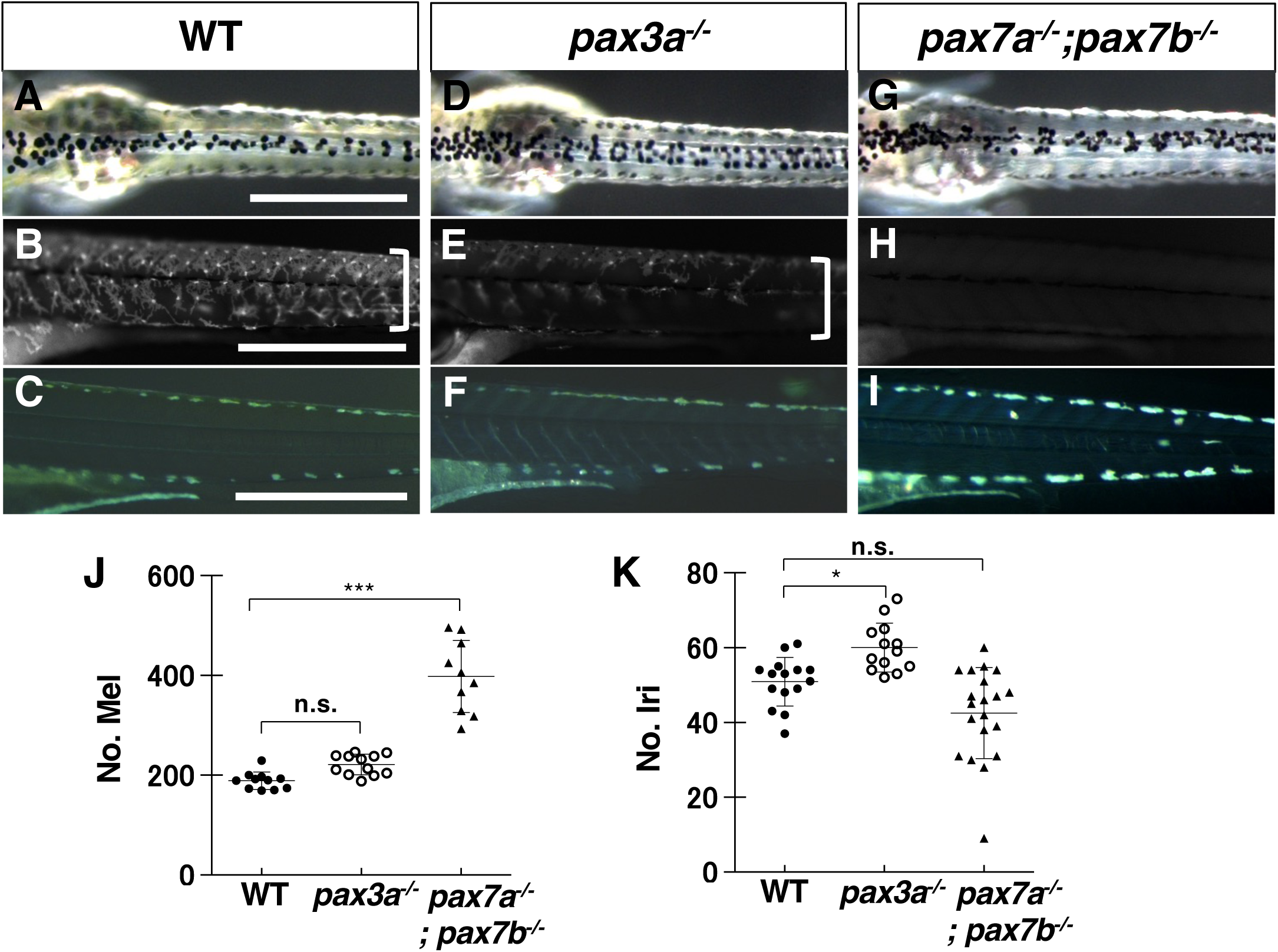
Phenotypes of zebrafish *pax3* and *pax7* mutants. (A-I) Zebrafish 4 dpf hatchlings. Pigment cell phenotypes are compared between WT (A, B, C), *pax3a*^-/-^ (D, E, F) and *pax7a^-/-^*; *pax7a^-/-^* (G, H, I). (A, D, G) Dorsal views of the trunk in dark field. (B, E, H) Lateral views of the trunk under UV light. (C, F, I) Lateral views of the trunk in dark field. (J, K) Cell counts. Melanophores (J) on the dorsal surface of the trunk are quantified. Iridophores in the dorsal and ventral edges of the trunk are quantified (K). In zebrafish, as in medaka, xanthophores are severely reduced in *pax3a*^-/-^ mutant hatchlings (xanthophores located in the yellowish part in A and D, and are more prominent in B and E, indicated by square brackets), and completely absent in *pax7a^-/-^*; *pax7a^-/-^* mutants (G, H), whereas melanophores are unaltered in *pax3a*^-/-^ mutants (A, D, J), while significantly increased in *pax7a^-/-^*; *pax7a^-/-^*mutants (G, J). Iridophores are slightly but significantly increased in *pax3a*^-/-^ mutants (C, F, K), while not changed in *pax7a^-/-^*; *pax7a^-/-^* mutants (I, K). Significant difference was determined by Kruscal-Wallis test. \*\*\**p*<0.05. n. s. = not significant. Scale bars = 250 µm

**Suppl. Fig. 6.**
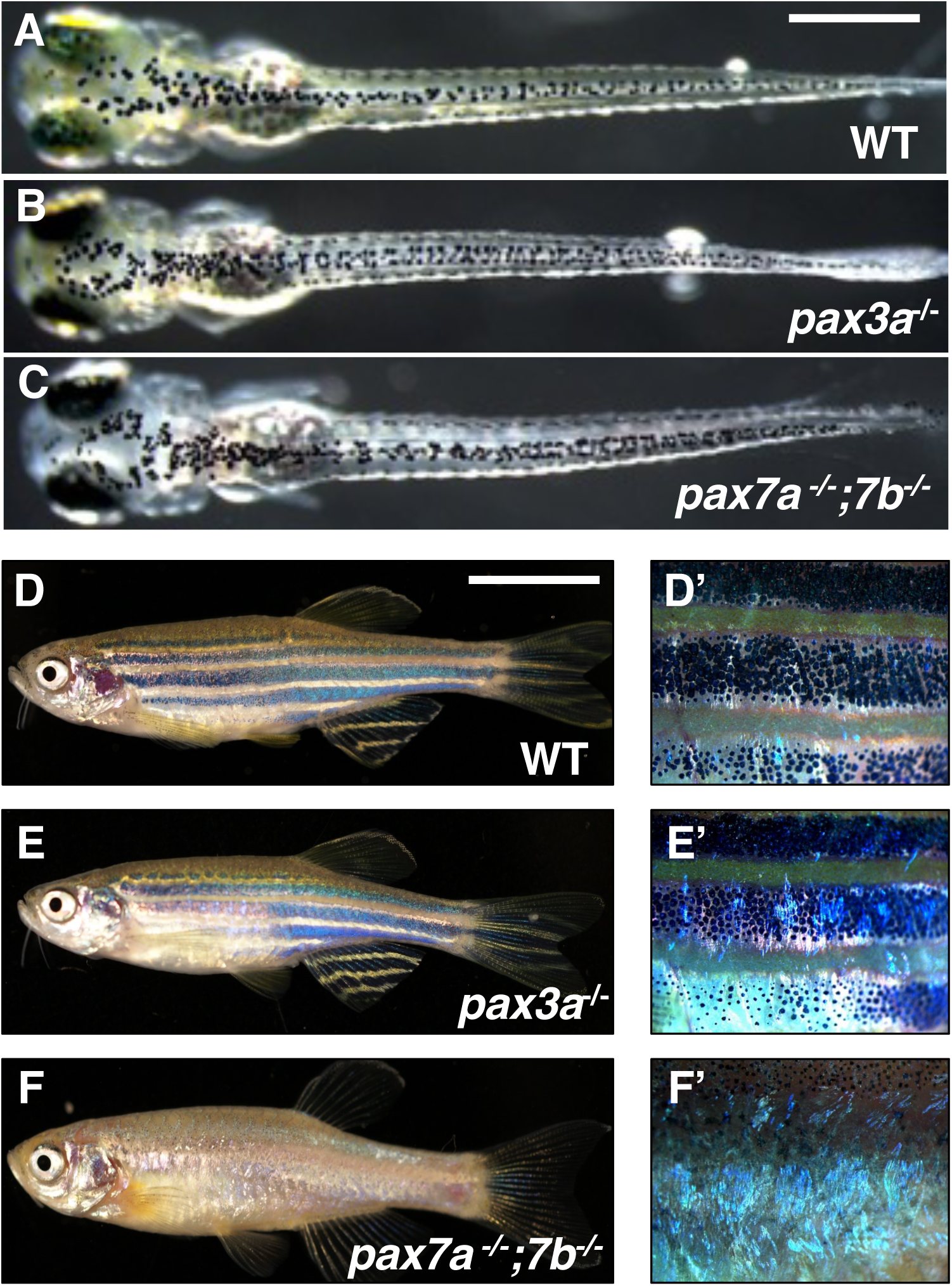
Larval and adult gross phenotypes of zebrafish *pax3a* and *pax7a; pax7b* mutants. (A-C) 4 dpf hatchlings. Dorsal views. Whole images of the same larvae shown in Suppl. Fig. 5. (D-F) 4-month-old adult fish. (A, D) Wild type (WT). (B, E) *pax3a^-/-^*. (C, F) *pax7a^-/-^; pax7b^-/-^*. (D’, E’, F’) Close-up images of the skin in D, E, F, respectively. Xanthophores are severely reduced in the *pax3a* mutant hatchling (A, B, see also Suppl. Fig. 5 A, B, D, E), but are restored in the adult (D, D’, E, E’). The *pax7a^-/-^; pax7b^-/-^* double mutant is completely defective for xanthophore formation not only at the hatching stage (C, see also Suppl. Fig. 5 G, H) but also at the adult stage (F, F’).

**Suppl. Fig. 7.**
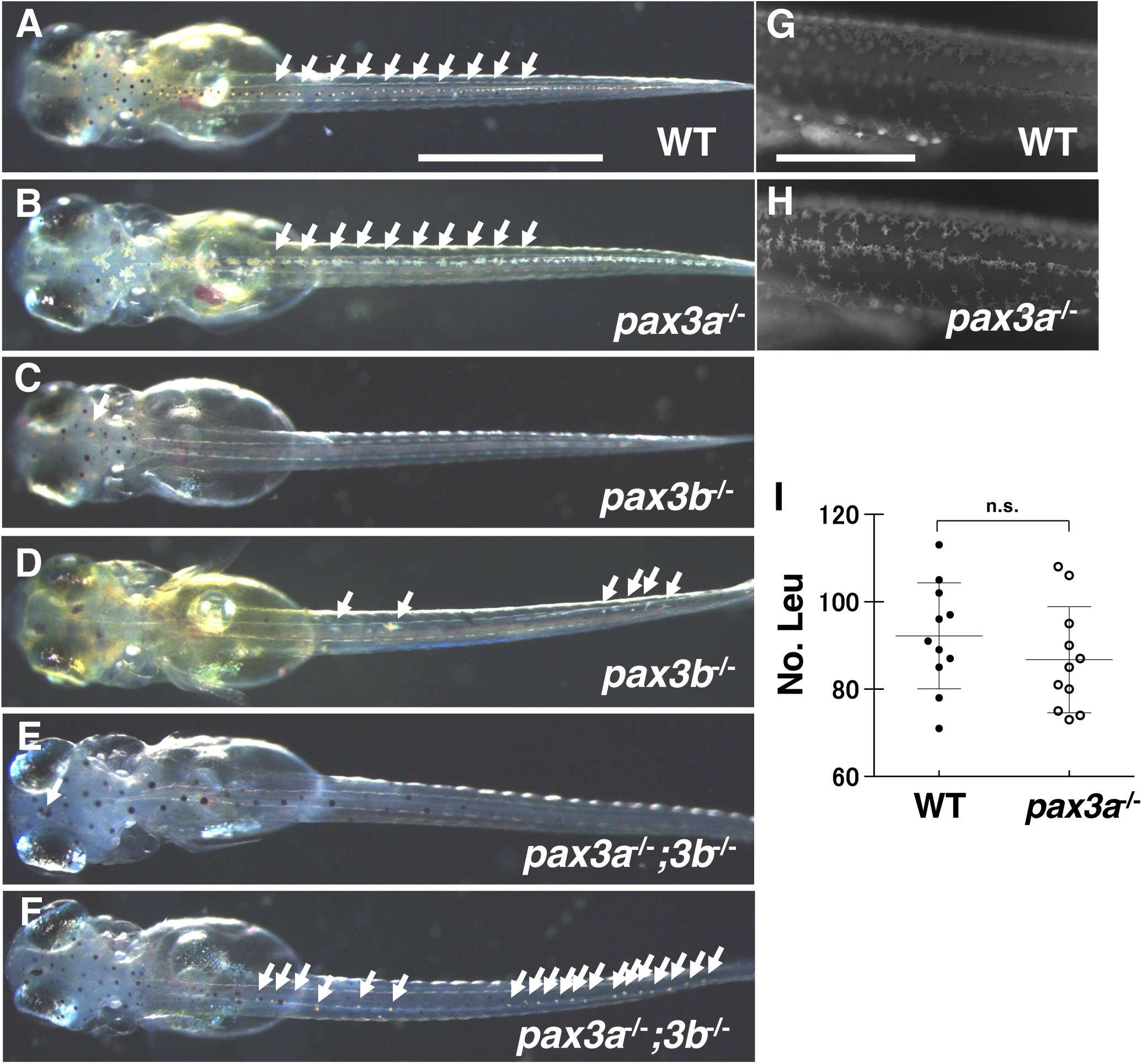
Medaka *pax3* mutant phenotypes. (A-F) 7 dpf hatchlings with d-rR background, in which melanophore pigmentation is severely defective. Dorsal views. Leucophores located in the dorsal midline were compared between WT (A) and *pax3a^-/-^* (B), *pax3b^-/-^*(C, D), *pax3a^-/-^; pax3b^-/-^* (E, F) mutants. The *pax3a* mutation does not appear to affect the pigment cell development, nor does it exacerbate the *pax3b* phenotype in the double homozygous *pax3a^-/-^; pax3b^-/-^* hatchling. (G, H) Autofluorescent xanthophores. Lateral views of the trunk under UV light. (I) Leucophore counts on the dorsal body in WT and *pax3a^-/-^* mutant. The double homozygotes failed to grow to adults. Xanthophore formation is not affected in *pax3a^-/-^* mutant (G, H). Arrows are pointing at leucophores on the body (A, B, D, F) and on the head (C, E). Scale bar = 0.5 mm (A-F) and 250 µm (G, H)

**Suppl. Fig. 8.**
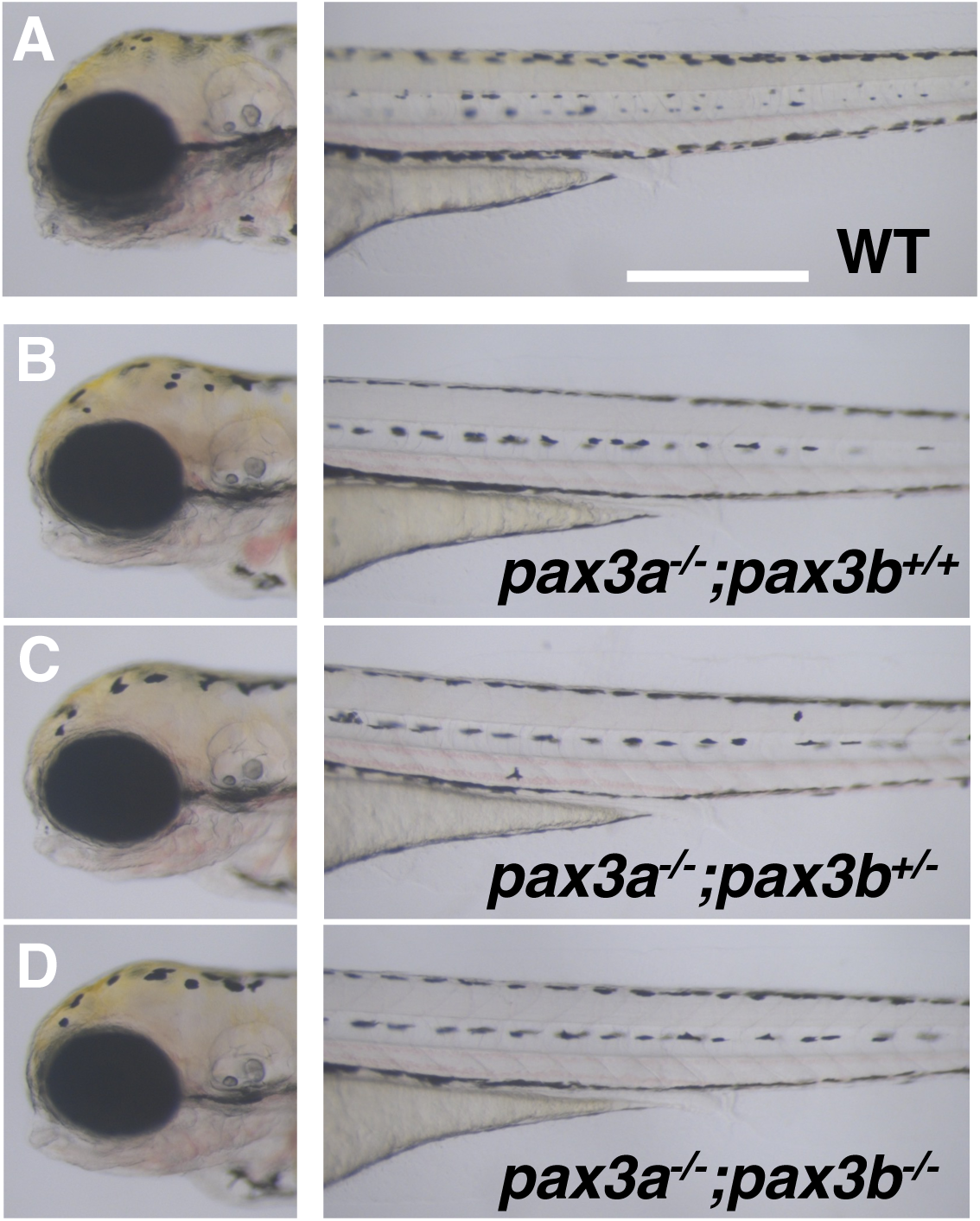
Zebrafish *pax3b* mutant phenotypes. (A-D) 4 dpf hatchlings. Lateral views. Left panels show head region. Right panels show trunk and tail regions. (A) WT. (B) *pax3a^-/-^*. (C) *pax3a^-/-^*, *pax3b^+/-^*. (D) *pax3a^-/-^; pax3b^-/-^*. Like medaka *pax3a* mutant, zebrafish *pax3b* homozygotes show no obvious pigment cell phenotypes (A, B). The *pax3b* mutation does not appear to exacerbate the *pax3a* phenotype in the *pax3a; pax3b* compound mutant hatchlings (C, D).

**Suppl. Fig. 9.**
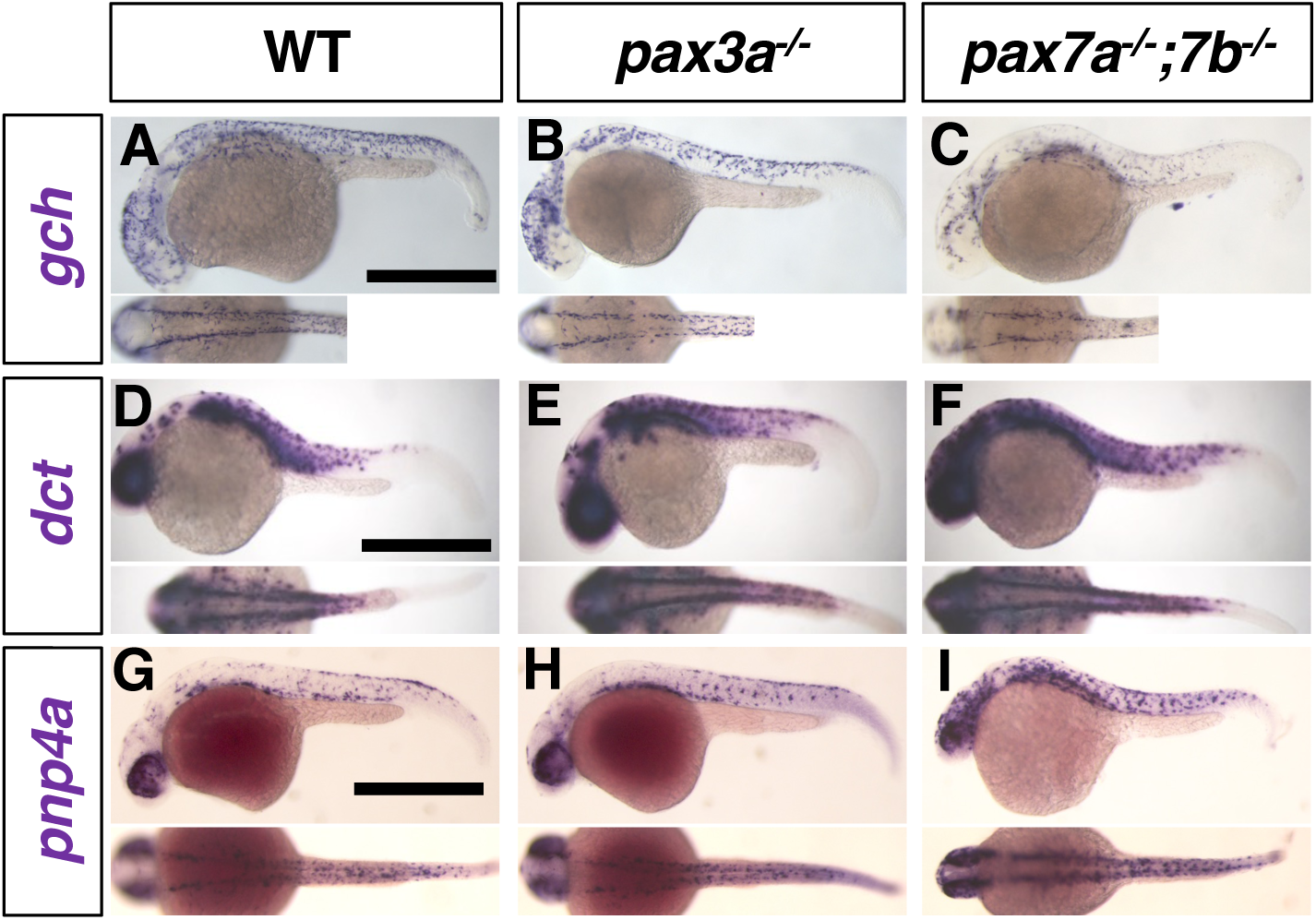
Pigment cell progenitors in zebrafish *pax3a* and *pax7a; pax7b* mutant embryos. 27 hpf. (A-C) *gch*. (D-F) *dct*. (G-I) *pnp4a*. (A, D, G) WT. (B, E, H) *pax3a^-/-^* mutant. (C, F, I) *pax7a^-/-^; pax7b^-/-^* mutant. (A-L) Lateral views at top and dorsal views at bottom. Scale bar = 250 µm The *gch*-expressing xanthophore progenitors are severely decreased in *pax3a* mutant (A, B) and in *pax7a; pax7b* mutant (C). The *dct-*expressing melanophore progenitors are comparable in WT (D) and in *pax3a* mutant €, while slightly increased in *pax7a; pax7b* mutant (F). Development of the *pnp4a*-expressing iridophore progenitors is not noticeably altered in the body in *pax3a* mutant (G, H), and unaltered or even increased in *pax7a; pax7b* mutant (I).

**Suppl. Fig. 10.**
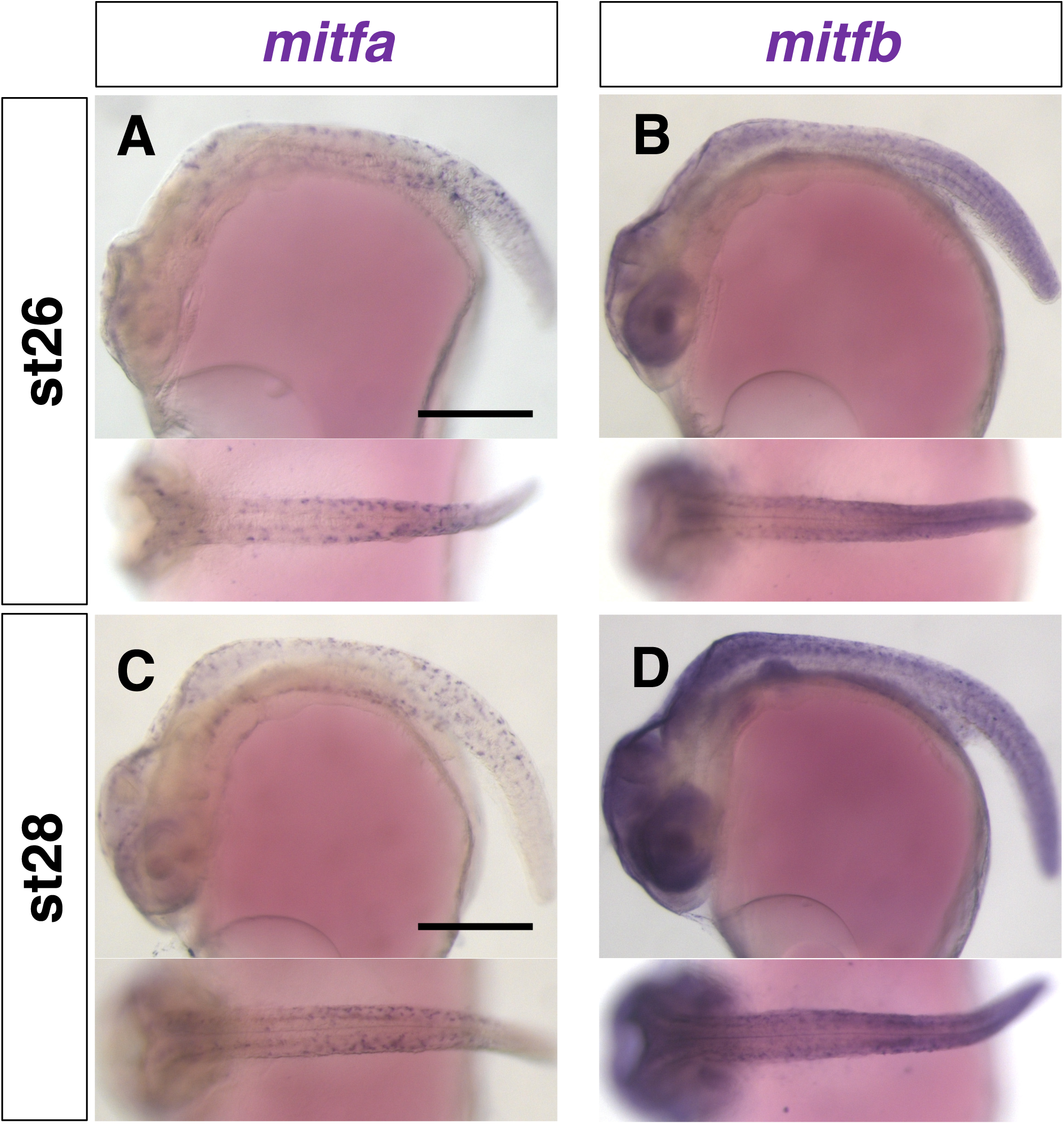
Comparison of *mitfa* and *mitfb* expression in medaka. (A, C,) *mitfa*. (B, D) *mitfb*. (A, B) Stage 26. (C, D) Stage 28. Lateral views top. Dorsal views below. All anterior to the left. Scale bars = 250 µm. In situ analyses were performed with medaka embryos. The *mitfa*-expressing cells are observed in the dorsal and lateral surface of the embryonic body and in the eye region (A, C). Stronger signals of the *mitfb*-expressing cells are detected at locations similar to those of the *mitfa*-expressing cells (B, D). The *mitfa* expression seems to be specific for the NCCs, perhaps pigment cell progenitors, whereas the *mitfb* expression is detected in the NCCs but also in the others, spinal cord, etc. Pigment cell progenitors with Mitf expression are likely to be detected more clearly with *mitfa* than with *mitfb* as a probe by in situ, which is the reason why we examined *mitfa* rather than *mitfb* in Fig. 5.

**Suppl. Fig. 11.**
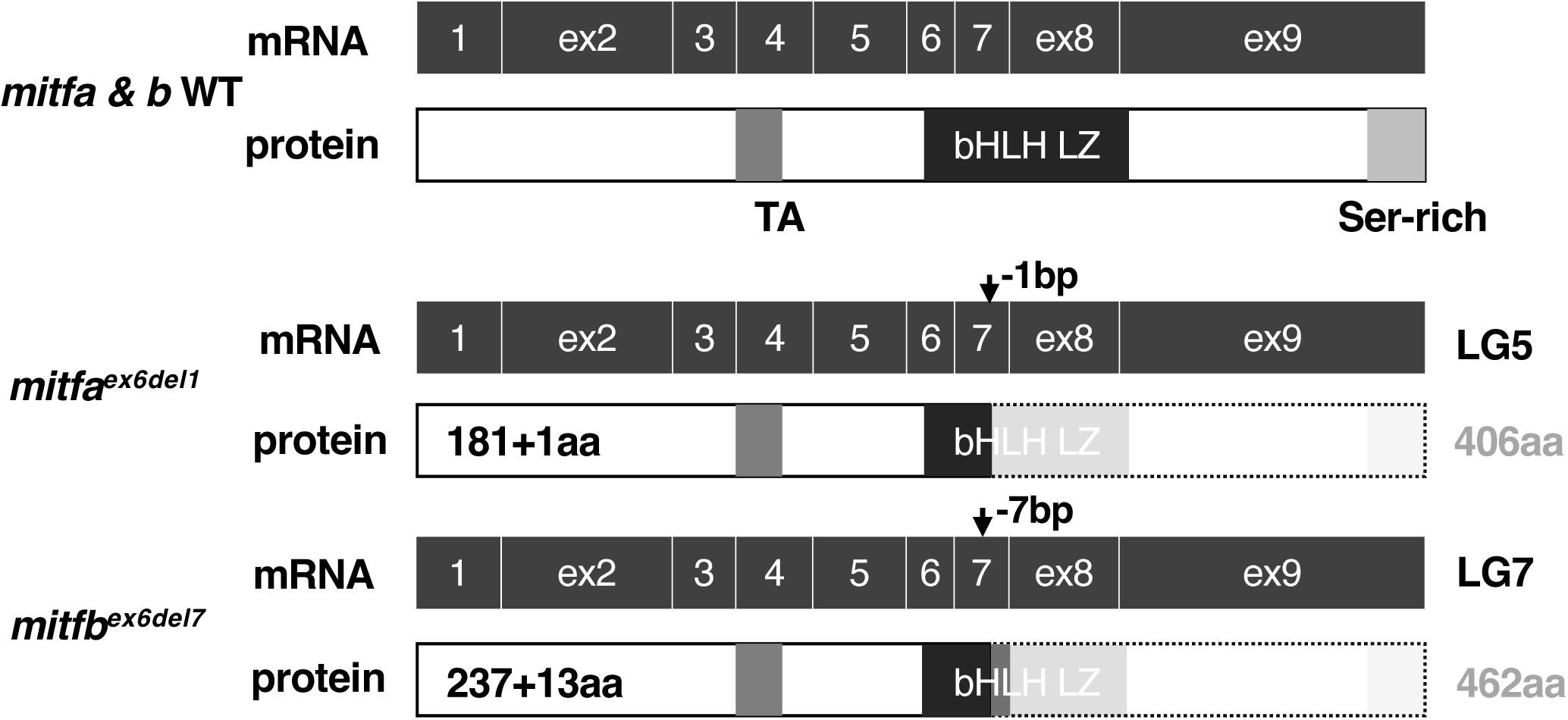
The *mitfa* and *mitfb* mutations in medaka. We generated mutants for *mitfa* and *mitfb* in medaka by CRISPR/Cas9. Both homozygotes were viable and fertile, and so we successfully established each strain. In addition, the *mitfa; mitfb* double mutants were also viable and fertile, and used for our analyses when necessary. Schematics of predicted primary structures are shown for wild type (WT) gene or mRNA and protein and mutant protein of Mitfa and Mitfb. The linkage group (LG) where the gene is located is shown on the right to the mRNA. The size (aa, amino acid) of the WT protein is shown right to the protein. The position and size (bp) of the deletion or insertion mutation is indicated by an arrow in the corresponding exon. The size of the mutant protein is given as the endogenous amino acid sequence + a de novo sequence following the frame-shift due to the deletion mutation (e.g., 181 + 1aa means that the Mitfa^ex6del1^ protein retains 181 N-terminal amino acids and one de novo amino acid). The Mitfb^ex6del7^ protein contains 237 N-terminal amino acids and 13 de novo amino acids. Neither of Mitfa and Mitfb proteins has a complete basic helix-loop-helix leucine zipper (bHLH-LZ) domain (black).

**Suppl. Fig. 12.**
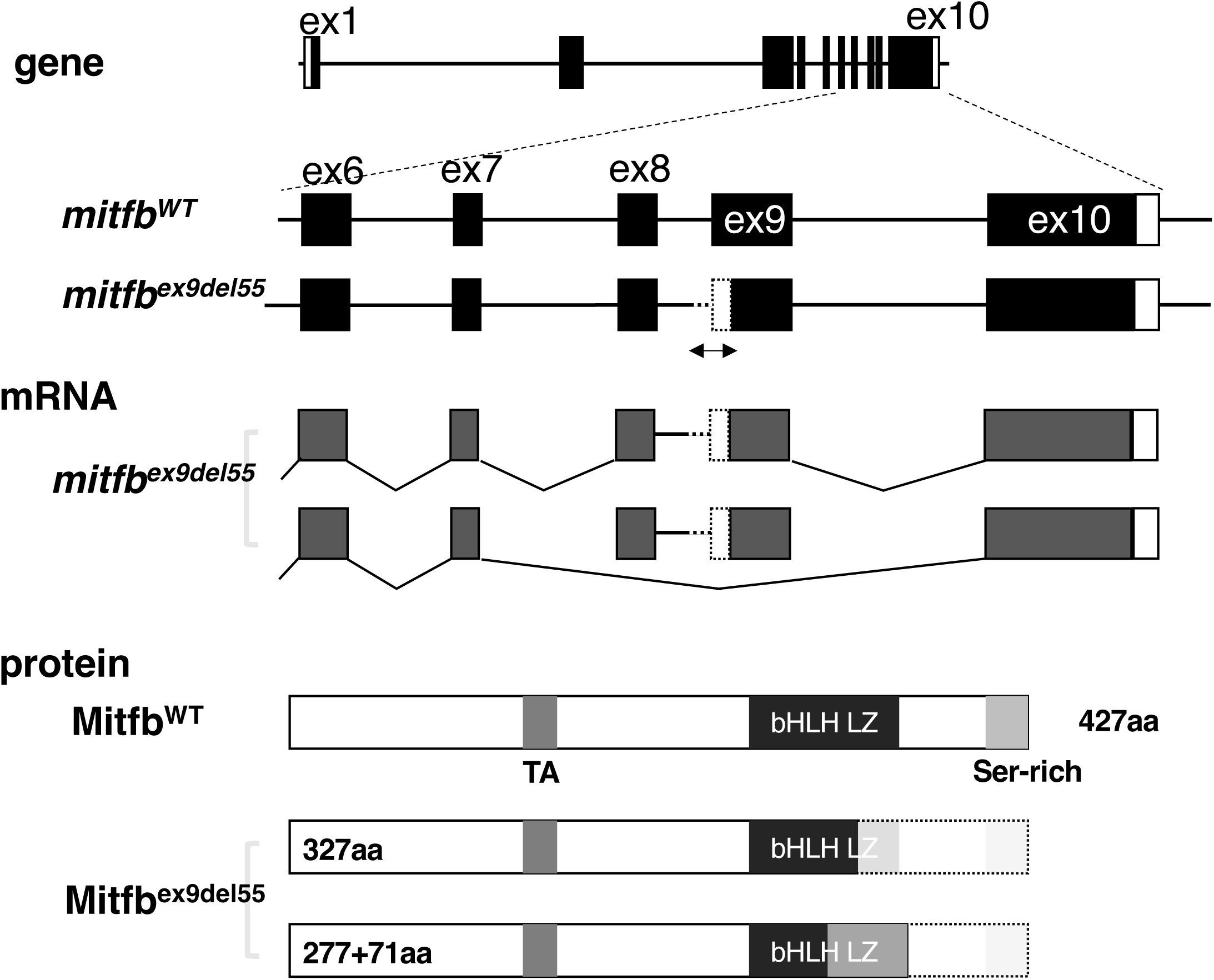
The *mitfb* mutation in zebrafish. In order to test if the requirement of Mitfs for xanthophore formation is conserved in zebrafish, we generated an *mitfb* mutation in zebrafish *mitfa^nacre^*background using CRISPR/Cas9. A guide RNA was designed to target exon 9: 5’– GAACAAGGGCACGATTCTGAAGG-3’ (Integrated DNA Technology) encoding the C-terminal half of bHLH domain. A 55 base pair deletion generated (*mitfb^ex9del55^*) expands the intron 8 and exon 9. Schematics of primary structures of the gene, mRNA and protein are shown for wild type (WT) and Mitfb^ex9del55^. RT-PCR and sequencing analyses revealed that the mutant allele produced two alternative mRNAs: one contained the intron 8 sequence between exon 8 and exon 9 but lacked the 55bp deletion sequence. The other mRNA skipped exon 8 and exon 9 with the 3’-end of exon 7 and the 5’-end of exon 10 joined. The mutant proteins have truncated C terminus of bHLH domain: the former mRNA containing intron 8 had a stop codon at the 5’-region of the intron, and the latter encoded de novo amino acid sequence (71 amino acids) after the absence of exon 8 and exon 9. None of the mutant proteins has a complete bHLH (dark gray) or a serine-rich domain (light gray).

**Suppl. Fig. 13.**
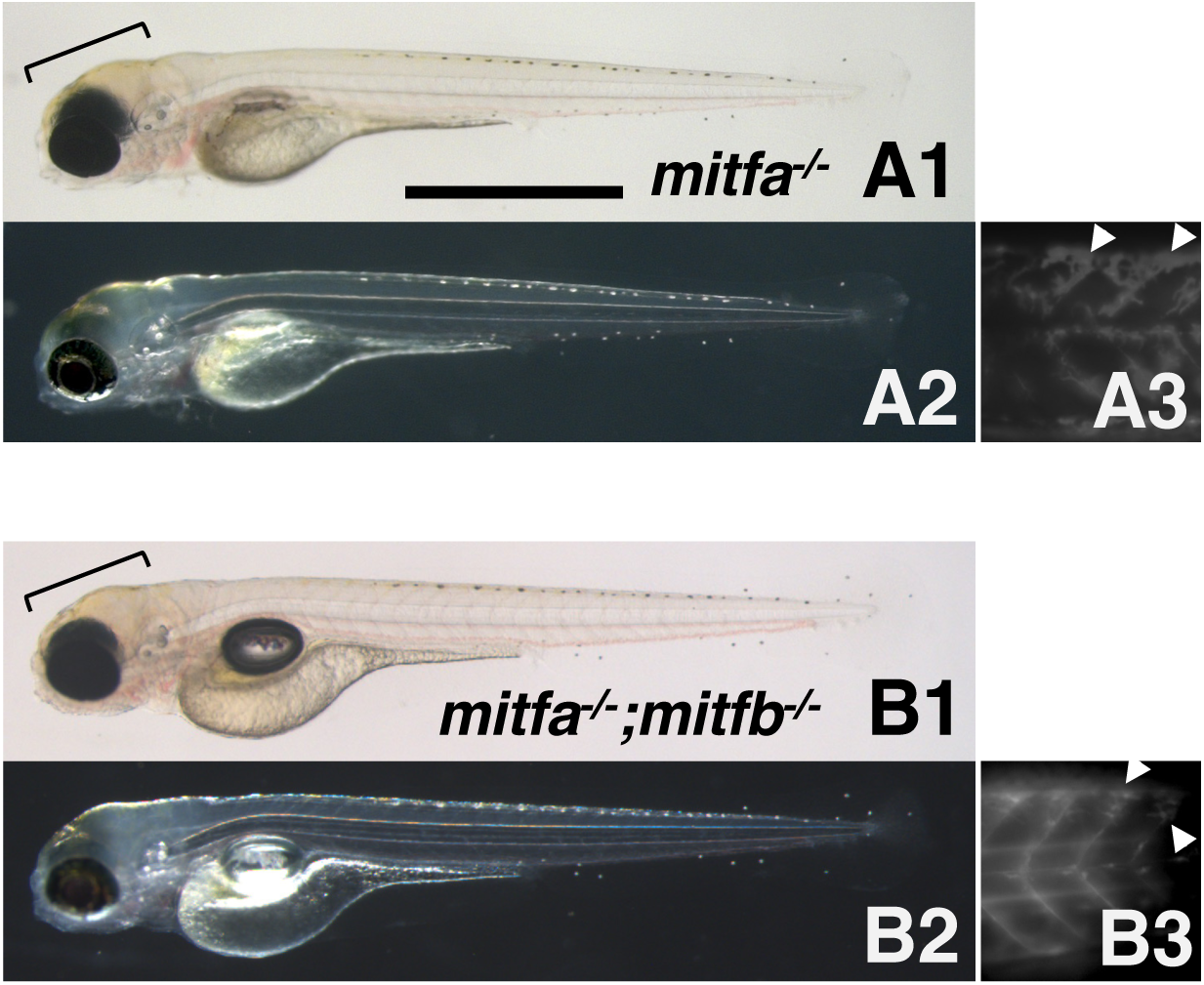
Phenotypes of zebrafish *mitfa* and *mitfb* double homozygotes. (A) 4 dpf *mitfa^-/-^* (*nacre*) hatchling. (B) 4 dpf *mitfa^-/-^*; *mitfb^-/-^* hatchling. (A1, B1) Normal transmission optics. (A2, B2) Dark-field epi-illumination optics. (A3, B3) Autofluorescence images showing xanthophores under UV light. Yellowish-pigmented xanthophores are observed in the dorsal head of the *mitfa^-/-^*hatchling (A1, A2). Autofluorescence emitted by xanthophores is clearly visible in the trunk of *mitfa^-/-^* (A3). Similarly, the *mitfa^-/-^*; *mitfb^-/-^* hatchling has xanthophores in the head (B1, B2) and autofluorescent cells in the trunk (B3). Brackets indicate pigmented xanthophores and arrowheads indicate autofluorescent (possibly immature) xanthophores. Scale bar = 1 mm Compared to a single *mitfa* mutant, *mitfa^-/-^*; *mitfb^-/-^* double mutants showed a slightly more severe reduction of xanthophores, with fewer fluorescent xanthophores on the trunk (A3, B3). However, unlike the medaka *mitfa*; *mitfb* double homozygotes, the zebrafish double mutants retained xanthophores on the head. Thus, the results suggest that in zebrafish xanthophore development may be partially dependent on Mitf function, but that some other factor(s) may compensate for the loss of Mitfs.

